# An experimentally determined evolutionary model dramatically improves phylogenetic fit

**DOI:** 10.1101/002899

**Authors:** Jesse D. Bloom

## Abstract

All modern approaches to molecular phylogenetics require a quantitative model for how genes evolve. Unfortunately, existing evolutionary models do not realistically represent the site-heterogeneous selection that governs actual sequence change. Attempts to remedy this problem have involved augmenting these models with a burgeoning number of free parameters. Here I demonstrate an alternative: experimental determination of a parameter-free evolutionary model via mutagenesis, functional selection, and deep sequencing. Using this strategy, I create an evolutionary model for influenza nucleoprotein that describes the gene phylogeny far better than existing models with dozens or even hundreds of free parameters. Emerging high-throughput experimental strategies such as the one employed here provide fundamentally new information that has the potential to transform the sensitivity of phylogenetic and genetic analyses.

## Introduction

The phylogenetic analysis of gene sequences is one of the most important and widely used computational techniques in all of biology. All modern phylogenetic algorithms require a quantitative evolutionary model that specifies the rate at which each site substitutes from one identity to another. These evolutionary models can be used to calculate the statistical likelihood of the sequences given a particular phylogenetic tree (Felsenstein, 1973). Phylogenetic relationships are typically inferred by finding the tree that maximizes this likelihood (Felsenstein, 1981) or by combining the likelihood with a prior to compute posterior probabilities of possible trees (Huelsenbeck et al., 2001).

Actual sequence evolution is governed by the rates at which mutations arise and the selection that subsequently acts upon them (Thorne et al., 2007; Halpern and Bruno, 1998). Unfortunately, neither of these aspects of the evolutionary process are traditionally known *a priori*. The standard approach in molecular phylogenetics is therefore to assume that sites evolve independently and identically, and then construct an evolutionary model that contains free parameters designed to represent features of mutation and selection (Goldman and Yang, 1994; Kosiol et al., 2007; Yang, 1994; Yang et al., 2000) This approach suffers from two major problems. First, although adding parameters enhances a model's fit to data, the parameter values must be estimated from the same sequences that are being analyzed phylogenetically – and so complex models can overfit the data (Posada and Buckley, 2004). Second, even complex models do not contain enough parameters to realistically represent selection, which is highly idiosyncratic to specific sites within a protein. Attempts to predict site-specific selection from protein structure have had limited success (Rodrigue et al., 2009; Kleinman et al., 2010), probably because even sophisticated computer programs cannot reliably predict the impact of mutations (Potapov et al., 2009).

Methods have been developed to infer site-specific selection from naturally occurring sequences (Rodrigue et al., 2010; Tamuri et al., 2012, 2014). Because the number of possible mutations is large, steps must be taken to ensure that these methods do not overfit the data (Rodrigue, 2013). But even when such steps are taken, the inferred site-specific selection parameters cannot easily be applied to phylogenetic analyses. The reason is that the selection parameters are generally inferred from the same naturally occurring sequences that are of phylogenetic interest – and parameters inferred from a data set cannot be used to analyze that same data set without additional procedures to avoid overfitting. The procedures that have been devised to restrain this problem of proliferating free parameters are complex, and generally require assuming that sites fall into only a limited number of different classes (Lartillot and Philippe, 2004; Le et al., 2008;Wu et al., 2013;Wang et al., 2008). Therefore, estimating site-specific selection from natural sequences is an imperfect method for inferring realistic evolutionary models for phylogenetic analyses.

Here I demonstrate a radically different approach for constructing quantitative evolutionary models: direct experimental measurement. This approach bypasses the aforementioned problem of proliferating free parameters because site-specific selection is measured experimentally without consideration of naturally occurring sequences. The evolutionary models constructed from these experiments therefore do not contain any parameters that must be estimated from the natural sequences that are being analyzed phylogenetically.

Specifically, using influenza nucleoprotein (NP) as an example, I experimentally estimate mutation rates via limiting-dilution passage and site-specific selection via deep mutational scanning (Fowler et al., 2010; Araya and Fowler, 2011), a combination of high-throughput mutagenesis, functional selection, and deep sequencing. I then show that these experimental measurements can be used to create a parameter-free evolutionary model describes the NP gene phylogeny far better than existing models with numerous free parameters. Finally, I discuss how the increasing availability of data from high-throughput experimental strategies such as the one employed here has the potential to transform analyses of genetic data by augmenting generic statistical models of evolution with detailed molecular-level information.

## Results

### Components of an experimentally determined evolutionary model

A phylogenetic evolutionary model specifies the rate at which one genotype is replaced by another. These rates of genotype substitution are determined by the underlying rates at which new mutations arise and the subsequent selection that acts upon them (Thorne et al., 2007; Halpern and Bruno, 1998). A standard assumption in molecular phylogenetics is that the rate of genotype substitution can be decomposed into independent substitution rates at individual sites. Here I make this assumption at the level of codon sites, and use *P_r,xy_* to denote the rate that site *r* substitutes from codon *x* to *y* given that the identity is already *x*. I further assume that it is possible to decompose *P_r,xy_* as

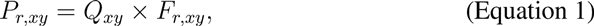

where *Q_xy_* is the rate of mutation from *x* to *y* (assumed to be constant across sites) and *F_r,xy_* is the site-specific probability that a mutation from *x* to *y* will fix at site *r* if it arises. Both are assumed to be constant over time.

Given the evolutionary model described by Equation 1, the challenge is to experimentally estimate the mutation rates *Q_xy_* and the fixation probabilities *F_r,xy_*. In the following sections, I describe these experiments.

### Measurement of mutation rates

A general challenge in quantifying mutation rates is the difficulty of separating mutations from the subsequent selection that acts upon them. To decouple mutation from selection, I utilized a previously described method for growing influenza viruses that package GFP in the PB1 segment (Bloom et al., 2010). The GFP does not contribute to viral growth and so is not under functional selection – therefore, substitutions in this gene accumulate at the mutation rate.

To drive the rapid accumulation of substitutions in the GFP gene, I performed limiting-dilution mutation-accumulation experiments (Halligan and Keightley, 2009). Specifically, I passaged 24 replicate populations of GFP-carrying influenza viruses by limiting dilution in tissue culture, at each passage serially diluting the virus to the lowest concentration capable of infecting target cells. Because each limiting dilution bottlenecks the population to one or a few infectious virions, mutations fix rapidly. After 25 rounds of passage, the GFP gene was Sanger sequenced for each replicate to identify 24 substitutions (Table 1, Table 2), for an overall rate of 5.6 × 10^−5^ mutations per nucleotide per tissue-culture generation – a value similar to that estimated previously by others using a somewhat different experimental approach (Parvin et al., 1986). The rates of different types of mutations are in Table 3, and possess expected features such as an elevation of transitions over transversions.

**Table 1:**
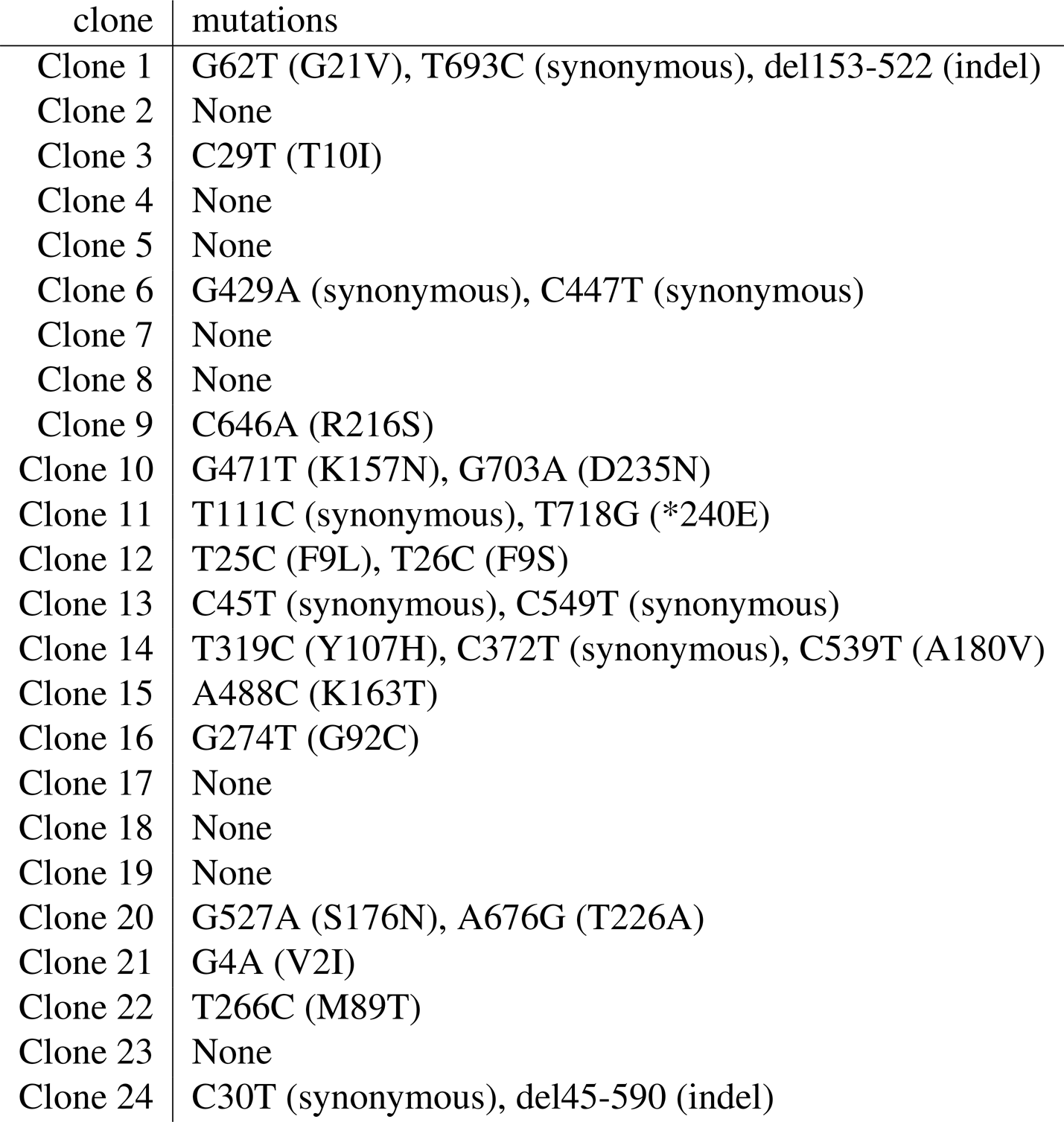
Mutations identified by sequencing the 720 nucleotide GFP gene packaged in the PB1 segment after 25 limiting-dilution passages for 24 independent replicates. The numbering is sequential beginning with the first nucleotide of the GFP start codon. For nonsynonymous mutations, the induced amino-acid change is indicated in parentheses.

| clone | mutations |
| --- | --- |
| Clone 1 | G62T (G21V), T693C (synonymous), del153-522 (indel) |
| Clone 2 | None |
| Clone 3 | C29T (T10I) |
| Clone 4 | None |
| Clone 5 | None |
| Clone 6 | G429A (synonymous), C447T (synonymous) |
| Clone 7 | None |
| Clone 8 | None |
| Clone 9 | C646A (R216S) |
| Clone 10 | G471T (K157N), G703A (D235N) |
| Clone 11 | T111C (synonymous), T718G (*240E) |
| Clone 12 | T25C (F9L), T26C (F9S) |
| Clone 13 | C45T (synonymous), C549T (synonymous) |
| Clone 14 | T319C (Y107H), C372T (synonymous), C539T (A180V) |
| Clone 15 | A488C (K163T) |
| Clone 16 | G274T (G92C) |
| Clone 17 | None |
| Clone 18 | None |
| Clone 19 | None |
| Clone 20 | G527A (S176N), A676G (T226A) |
| Clone 21 | G4A (V2I) |
| Clone 22 | T266C (M89T) |
| Clone 23 | None |
| Clone 24 | C30T (synonymous), del45-590 (indel) |

**Table 2:**
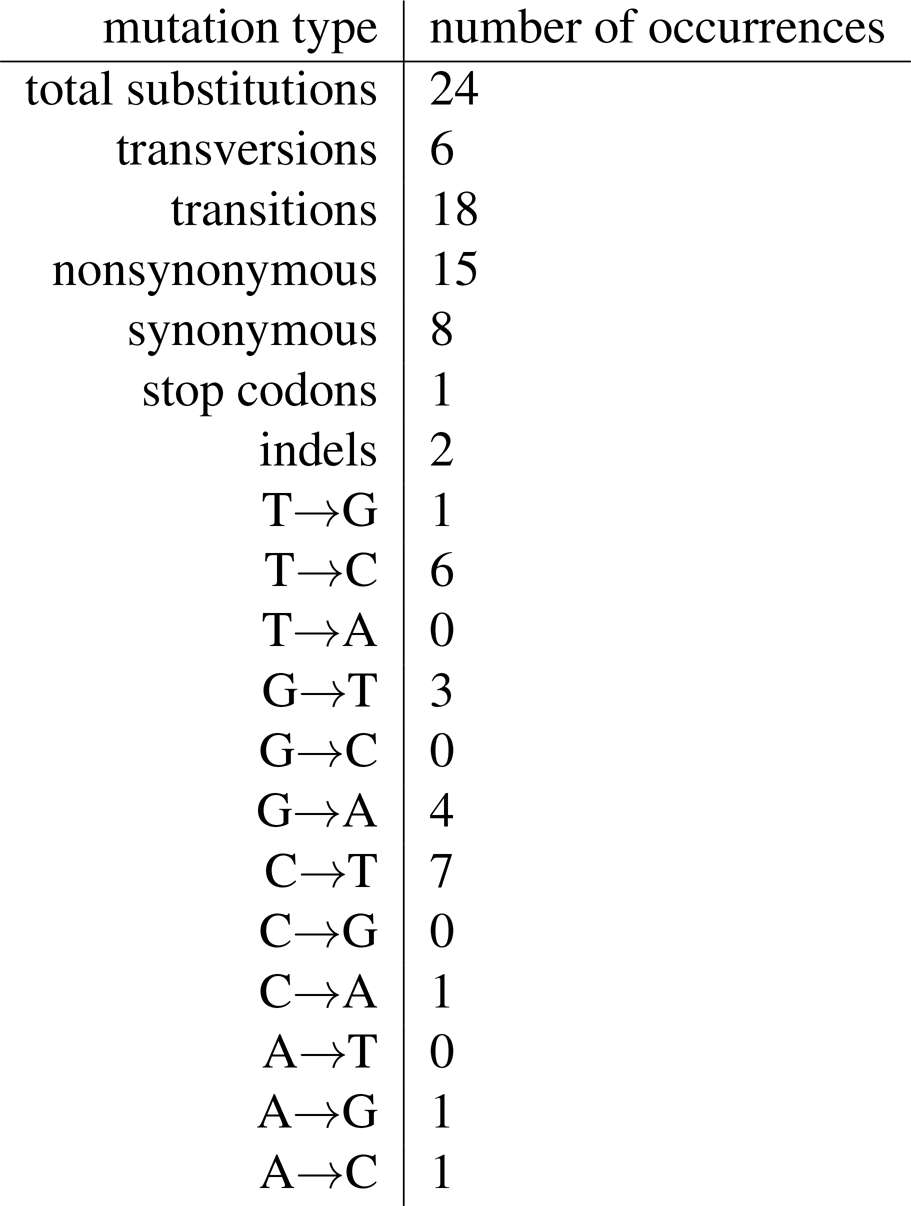
Counts for different types of mutations after the 25 limiting-dilution passages. The numbers are calculated from Table 1. Given that GFP is 720 nucleotides long, the data suggest a viral mutation rate of 5.6 × 10^−5^ mutations per nucleotide per tissue-culture generation.

**Table 3:** Influenza mutation rates. Numbers represent the probability a site that has the parent identity will mutate to the specified nucleotide in a single tissue-culture generation, and are calculated from Table 1 and Table 2 after adding one pseudocount to each mutation type. Mutations are in pairs because an observed change of A → G can derive either from this mutation on the sequenced strand or a T → C on the complementary strand, and so the paired mutations are indistinguishable assuming that the same mutational process applies to both strands of the replicated nucleic acid molecule. The numbers are the estimated rates of each individual mutation, so for example the observed rate of change from A→G is 2 × (2:4 × 10^−5^) since this change can arise from either of the two mutations A→G and T→C.

| mutation type | rate |
| --- | --- |
| A→G, T→C (transition) | $2.4 \times 10^{-5}$ |
| G→A, C→T (transition) | $2.3 \times 10^{-5}$ |
| A→C, T→G (transversion) | $9.0 \times 10^{-6}$ |
| C→A, G→T (transversion) | $9.4 \times 10^{-6}$ |
| A→T, T→A (transversion) | $3.0 \times 10^{-6}$ |
| G→C, C→G (transversion) | $1.9 \times 10^{-6}$ |

Because an observed mutation of A → G can arise from either this change on the sequenced strand or a change of T → C on the complementary strand, then assuming that the same molecular mutation process affects both strands, there are only the six different mutation rates shown in Table 3. Specifically, let *R_m→n_* represent the rate at which nucleotide *m* mutates to *n* given that the identity is already *m*, and let *m_c_* denote the complement of *m* (so for example A*_c_* is T). The assumption that the same molecular mutation process affects both strands means that 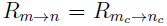. An additional empirical observation from Table 3 is that the mutation rates for influenza are approximately symmetric, with the rate of each mutation approximately equal to its reversal 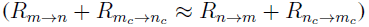. Because it somewhat simplifies computational aspects of the subsequent phylogenetic analyses, I enforce this empirical observation of approximately symmetric mutation rates to be exactly true by taking the rates of mutations and their reversals to be the average of the two. With the further assumption that codon mutations occur a single nucleotide at a time, the mutation rates *Q_xy_* from codon *x* to *y* are estimated from the experimental data in Table 3 as

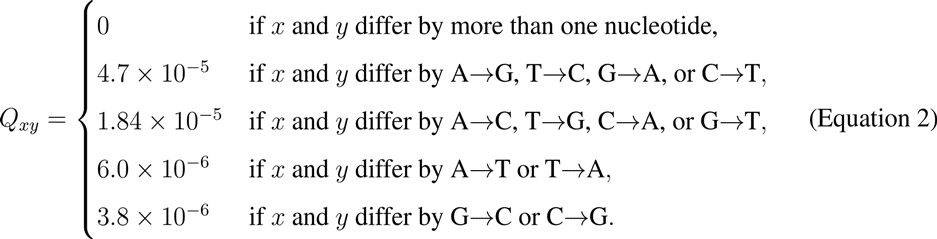

These mutation rates define the first term in the evolutionary model specified by Equation 1.

### Deep mutational scanning to assess effects of mutations on NP

Estimation of the fixation probabilities *F_r,xy_* in Equation 1 requires quantifying the effects of all ≈ 10^4^ possible amino-acid mutations to NP. Such large-scale assessments of mutational effects are feasible with the advent of deep mutational scanning, a recently developed experimental strategy of high-throughput mutagenesis, selection, and deep sequencing (Fowler et al., 2010; Araya and Fowler, 2011) that has now been applied to several genes (Fowler et al., 2010; Melamed et al., 2013; Traxlmayr et al., 2012; Starita et al., 2013; Roscoe et al., 2013). Applying this experimental strategy to NP requires creating large libraries of random gene mutants, using these genes to generate pools of mutant influenza viruses which are then passaged at low multiplicity of infection to select for functional variants, and finally using Illumina sequencing to assess the frequency of each mutation in the input mutant genes and the resulting viruses. Mutations that interfere with NP function or stability impair or ablate viral growth, since NP plays an essential role in influenza genome packaging, replication, and transcription (Ye et al., 2006; Portela and Digard, 2002). Such mutations will therefore be depleted in the mutant viruses relative to the input mutant genes.

Most previous applications of deep mutational scanning have examined single-nucleotide mutations to genes, since such mutations can easily be generated by error-prone PCR or other nucleotide-level mutagenesis techniques. However, many amino-acid mutations are not accessible by single-nucleotide changes. I therefore used a PCR-based strategy to construct codon-mutant libraries that contained multi-nucleotide (i.e. GGC → <u>ACT</u>) as well as single-nucleotide (i.e. GGC → <u>A</u>GC) mutations. The use of codon-mutant libraries has an added benefit during the subsequent analysis of the deep sequencing when trying to separate true mutations from errors, since the majority (54 of 63) possible codon mutations involve multi-nucleotide changes whereas sequencing and PCR errors generate almost exclusively single-nucleotide changes. I used identical experimental procedures to construct two codon-mutant libraries of NP from the wild type (WT) human H3N2 strain A/Aichi/2/1968, and two from a variant of this NP with a single amino-acid substitution (N334H) that enhances protein stability (Gong et al., 2013; Ashenberg et al., 2013). These codon-mutant libraries are termed WT-1, WT-2, N334H-1, and N334H-2. Each of these four mutant libraries contained *>* 10^6^ unique plasmid clones. Sanger sequencing of 30 clones drawn roughly equally from the four libraries revealed that the number of codon mutations per clone followed a Poisson distribution with a mean of 2.7 (Figure 1). These codon mutations were distributed roughly uniformly along the gene sequence and showed no obvious biases towards specific mutations (Figure 1). Most of the ≈ 10^4^ unique amino-acid mutations to NP therefore occur in numerous different clones in the four libraries, both individually and in combination with other mutations.

**Figure 1:**
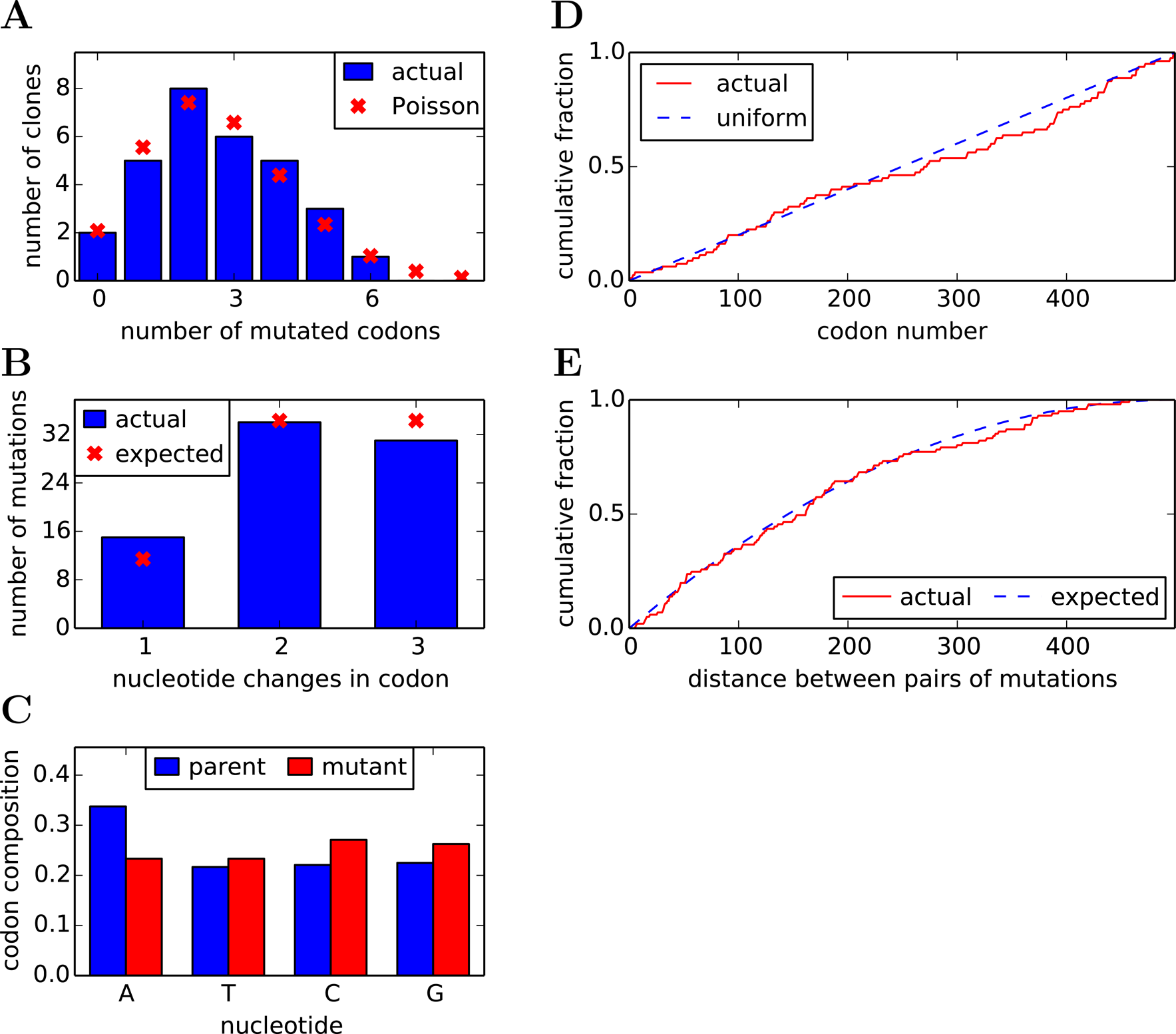
The codon-mutant libraries as assessed by Sanger sequencing 30 individual clones. **(A)** The clones have an average of 2.7 codon mutations and 0.1 indels per full-length NP coding sequence, with the number of mutated codons per gene following an approximately a Poisson distribution. **(B)** The number of nucleotide changes per codon mutation is roughly as expected if each codon is randomly mutated to any of the other 63 codons, with a slight elevation in single-nucleotide mutations. **(C)** The mutant codons have a uniform base composition. **(D)** Mutations occur uniformly along the primary sequence. **(E)** In clones with multiple mutations, there is no tendency for mutations to cluster. Shown is the actual distribution of pairwise distances between mutations in all multiply mutated clones compared to the distribution generated by 1,000 simulations where mutations are placed randomly along the primary sequence of each multiple-mutant clone. The data and code for this figure are available at https://github.com/jbloom/SangerMutantLibraryAnalysis/tree/v0.21.

To assess effects of the mutations on viral replication, the plasmid mutant libraries were used to create pools of mutant influenza viruses by reverse genetics (Hoffmann et al., 2000). The viruses were passaged twice in tissue culture at low multiplicity of infection to enforce a linkage between genotype and phenotype. The NP gene was reverse-transcribed and PCR-amplified from viral RNA after each passage, and similar PCR amplicons were generated from the plasmid mutant libraries and a variety of controls designed to quantify errors associated with sequencing, reverse transcription, and viral passage (Figure 2). The entire process outlined in Figure 2 was performed in parallel but separately for each of the four mutant libraries (WT-1, WT-2, N334H-1, and N334H-2) in what will be termed one experimental *replicate*. This entire process of viral creation, passaging, and sequencing was then repeated independently for all four libraries in a second experimental replicate. The two independent replicates will be termed *replicate A* and *replicate B*.

**Figure 2:**
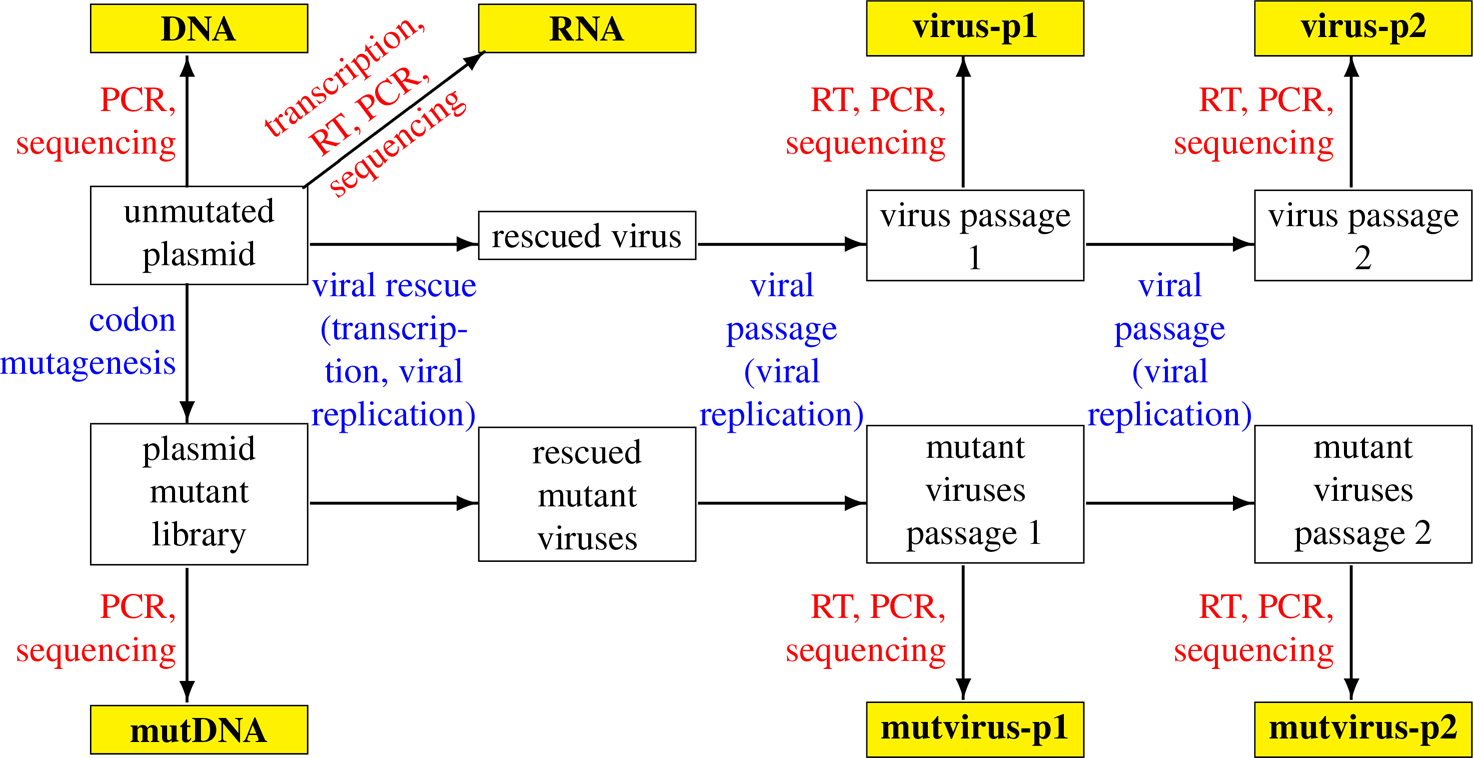
Design of the deep mutational scanning experiment. The sequenced samples are in yellow. Blue text indicates sources of mutation and selection; red text indicates sources of errors. The comparison of interest is between the mutation frequencies in the **mutDNA** and **mutvirus** samples, since changes in frequencies between these samples represent the action of selection. However, since some of the experimental techniques have the potential to introduce errors, the other samples are also sequenced to quantify these unintended sources of error. Each of the two experimental replicates (*replicate A* and *replicate B*) involved independently repeating the entire viral rescue, viral passaging, and sequencing process for each of the four plasmid mutant libraries (WT-1, WT-2, N334H-1, and N334H-2).

The mutation frequencies in all samples were quantified by Illumina sequencing, using over-lapping paired-end reads to reduce errors (Supplementary figure 1). Each sample produced ≈ 10^7^ paired reads that could be aligned to NP, providing an average of ≈ 5 × 10^5^ calls per codon (Supplementary figure 2). Sequencing of unmutated NP plasmid revealed a low rate of errors, which were almost exclusively single-nucleotide changes (Figure 3). As expected, the plasmid mutant libraries contained a high frequency of single and multi-nucleotide codon mutations (Figure 3). Mutation frequencies for unmutated RNA or viruses created from unmutated NP plasmid were only slightly above the sequencing error rate (Figure 3), indicating that reverse-transcription and viral replication introduced few mutations relative to the targeted mutagenesis in the plasmid libraries. Mutation frequencies were reduced in the mutant viruses relative to the mutant plasmids used to create these viruses, particularly for nonsynonymous and stop-codon mutations (Figure 3) – consistent with selection purging deleterious mutations. These results indicate that the deep mutational scanning experiment successfully introduced many of the NP variants in the plasmid mutant libraries into mutant viruses which were then subjected to purifying selection against mutations that interfered with viral replication.

**Figure 3:**
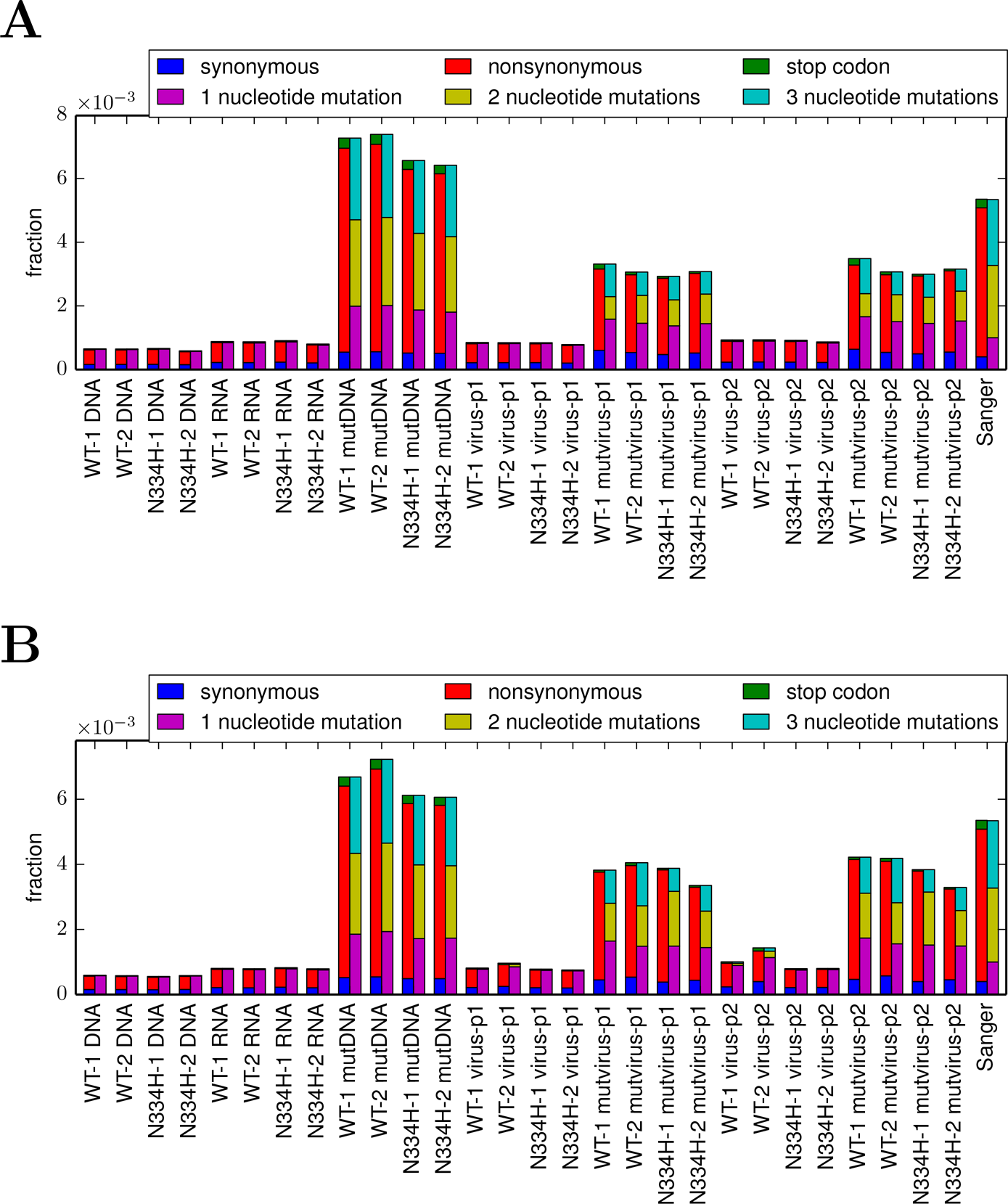
Per-codon mutation frequencies for each library (WT-1, WT-2, N334H-1, N334H-2) in **(A)** *replicate A* or **(B)** *replicate B*. The samples are named as in Figure 2. Errors due to Illumina sequencing (**DNA** sample), reverse-transcription (**RNA** sample), and viral replication (**virus-p1** and **virus-p2** samples) are rare, and are mostly single-nucleotide changes. The codon-mutant libraries (**mutDNA**) contain a high frequency of single- and multi-nucleotide changes as expected from Sanger sequencing (rightmost bars of this plot and Figure 1; note that Sanger sequencing is not subject to Illumina sequencing errors that affect all other samples). Mutations are reduced in **mutvirus** samples relative to **mutDNA** plasmids used to create these mutant viruses, with most of the reduction in stop-codon and nonsynonymous mutations – as expected if deleterious mutations are purged by purifying selection. Details of the analysis used to generate these figures are at http://jbloom.github.io/mapmuts/example_2013Analysis_Influenza_NP_Aichi68.html.

**Figure 4:**
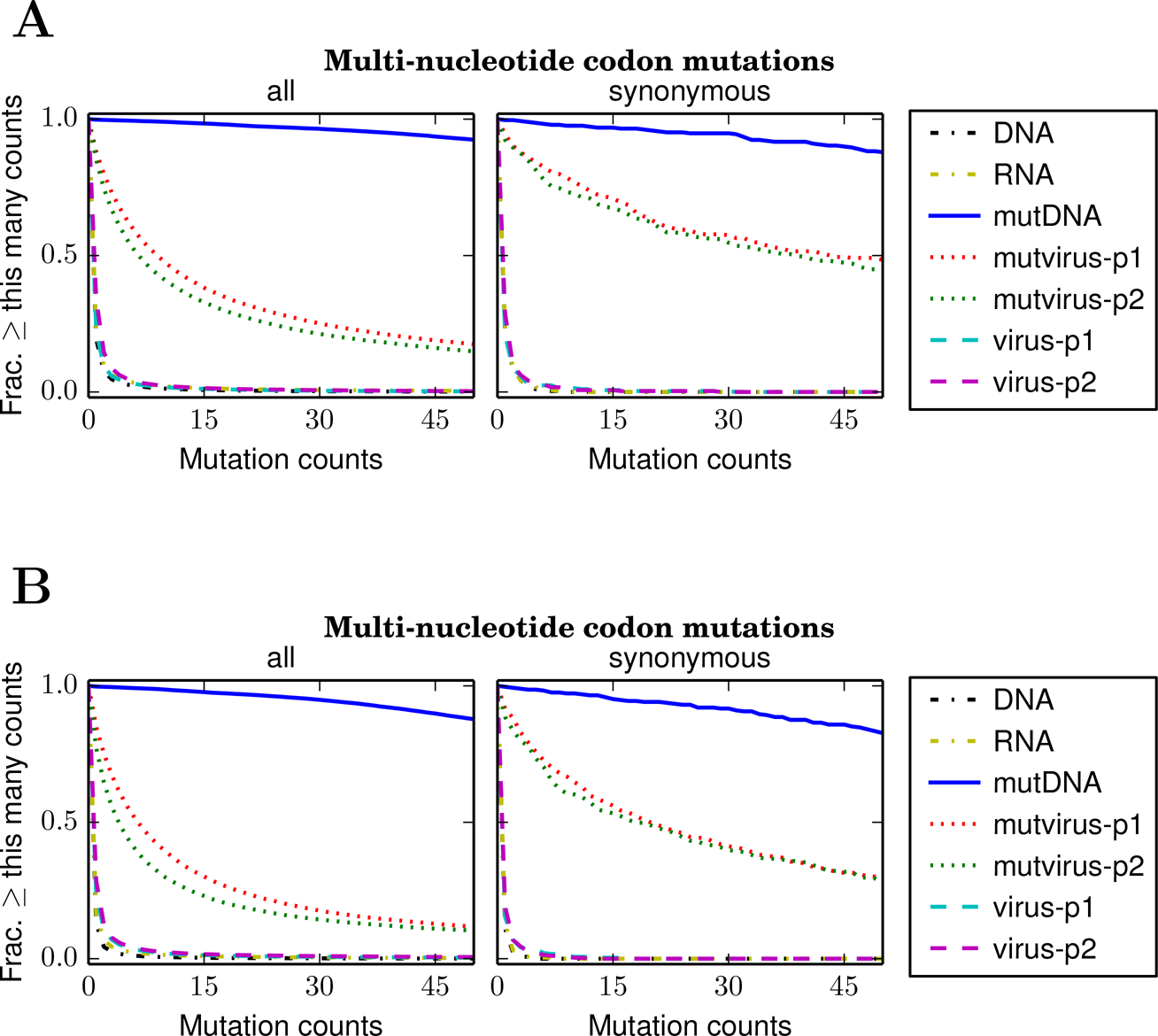
The completeness with which mutations were sampled in the mutant plasmids and viruses, as assessed by the counts for each multi-nucleotide codon mutation in the combined libraries of **(A)** *replicate A* or **(B)** *replicate B*. Restricting these plots to multi-nucleotide codon mutations avoids confounding effects from sequencing errors, which typically generate single-nucleotide codon mutations. Very few multi-nucleotide codon mutations are observed more than once in the unmutagenized controls (**DNA, RNA, virus-p1, virus-p2**). Nearly all multi-nucleotide codon mutations are observed many times in the mutant plasmid libraries (**mutDNA**). About half the multi-nucleotide codon mutations are found at least five times in the mutant viruses (**mutvirus-p1, mutvirus-p2**), indicating that at least half the possible mutations were incorporated into a virus. However, this is only a lower bound, since deleterious mutations will be absent from the mutant viruses due to purifying selection. If the analysis is restricted to synonymous multi-nucleotide codon mutations (which are less likely to be deleterious), then over 75% of the possible mutations were incorporated into a virus. This is still only a lower bound, since even synonymous mutations are sometimes strongly deleterious to influenza (Marsh et al., 2008). The completeness with which amino-acid mutations are sampled is higher due to the redundancy of the genetic code. Note that *replicate A* is superior to *replicate B* in terms of the completeness with which the mutations are sampled by the mutant viruses. Details of the analysis used to generate these figures are at http://jbloom.github.io/mapmuts/example_2013Analysis_Influenza_NP_Aichi68.html.

**Figure 5:**
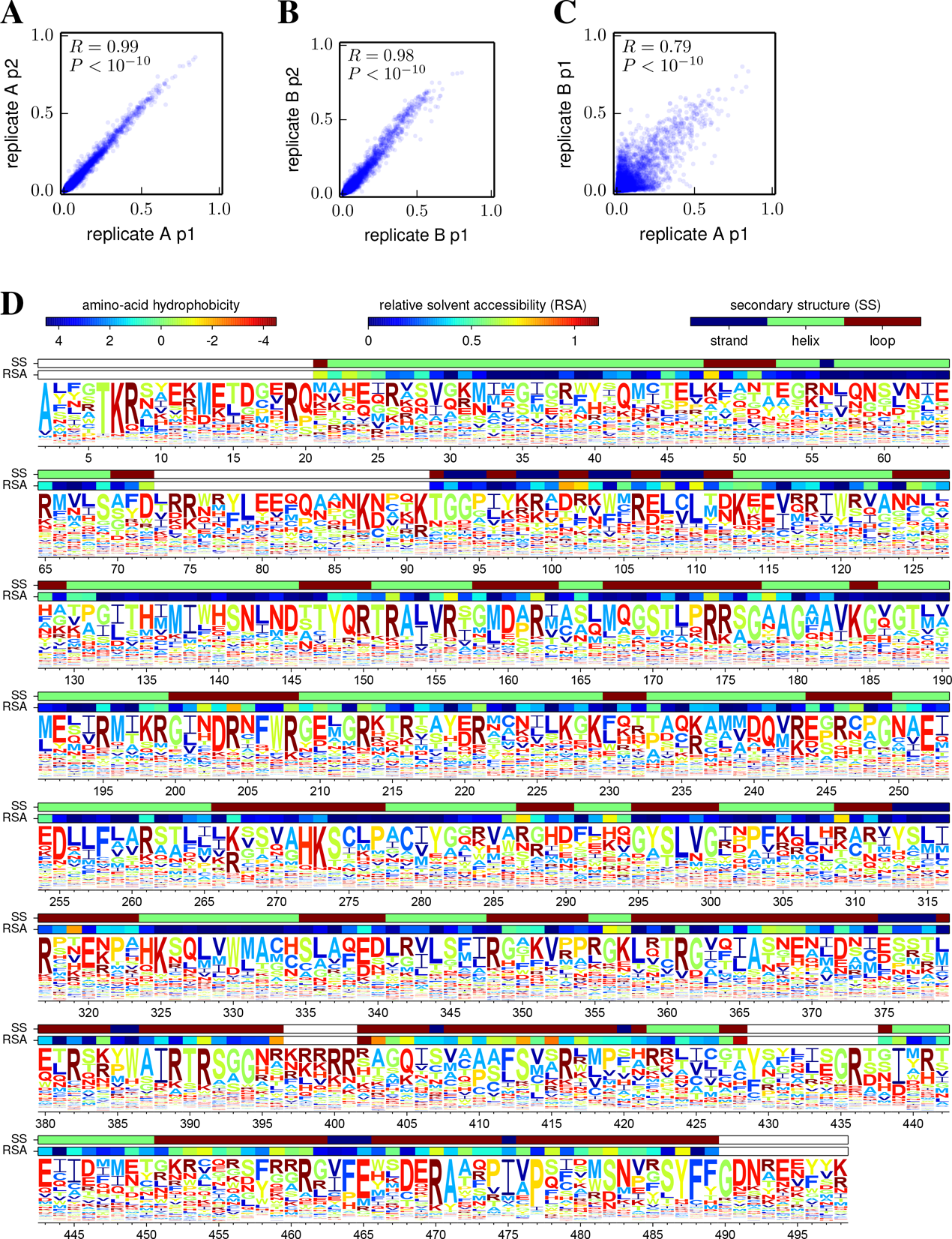
Amino-acid preferences. **(A), (B)** Preferences inferred from passage 1 and 2 are similar within each replicate, indicating that most selection occurs during initial viral creation and passage, and that technical variation is small. **(C)** Preferences from the two independent replicates are also correlated, but less perfectly. The increased variation is presumably due to stochasticity during the independent viral creation from plasmids for each replicate. **(D)** Preferences for all sites in NP (the N-terminal Met was not mutagenized) inferred from passage 1 of the combined replicates. Letters heights are proportional to the preference for that amino acid and are colored by hydrophobicity. Relative solvent accessibility and secondary structure are overlaid for residues in crystal structure. Correlation plots show Pearson’s *R* and *P*-value. Numerical data for **(D)** are in Supplementary file 1. The preferences are consistent with existing knowledge about mutations to NP (Table 4, Table 5). The computer code used to generate this figure is at http://jbloom.github.io/mapmuts/example_2013Analysis_Influenza_NP_Aichi68.html.

A key question is the extent to which the possible mutations were sampled in both the plasmid mutant libraries and the mutant viruses created from these plasmids. The deep mutational scanning would not achieve its goal if only a small fraction of possible mutations are sampled by the mutant plasmids or by the mutant viruses created from these plasmids (the latter might be the case if there is a bottleneck during virus creation such that all viruses are generated from only a few plasmids). Fortunately, Figure 4 shows that the sampling of mutations was quite extensive in both the mutant plasmids and the mutant viruses. Specifically, Figure 4 suggests that for each replicate, nearly all codon mutations were sampled numerous times in the plasmid mutant libraries, and that over 75% of codon mutations were sampled by the mutant viruses. Figure 4 also suggests that *replicate A* was technically superior to *replicate B* in the thoroughness with which mutations were sampled by the mutant viruses. Because most amino acids are encoded by multiple codons, the fraction of amino-acid mutations sampled in each replicate is even higher than the *>* 75% of sampled codon mutations. So while the experiments may not have exhaustively examined every possible codon mutation, the thoroughness of sampling is certainly sufficient to make the sort of statistical inferences about mutational effects that are necessary to construct a quantitative evolutionary model.

**Figure 6:**
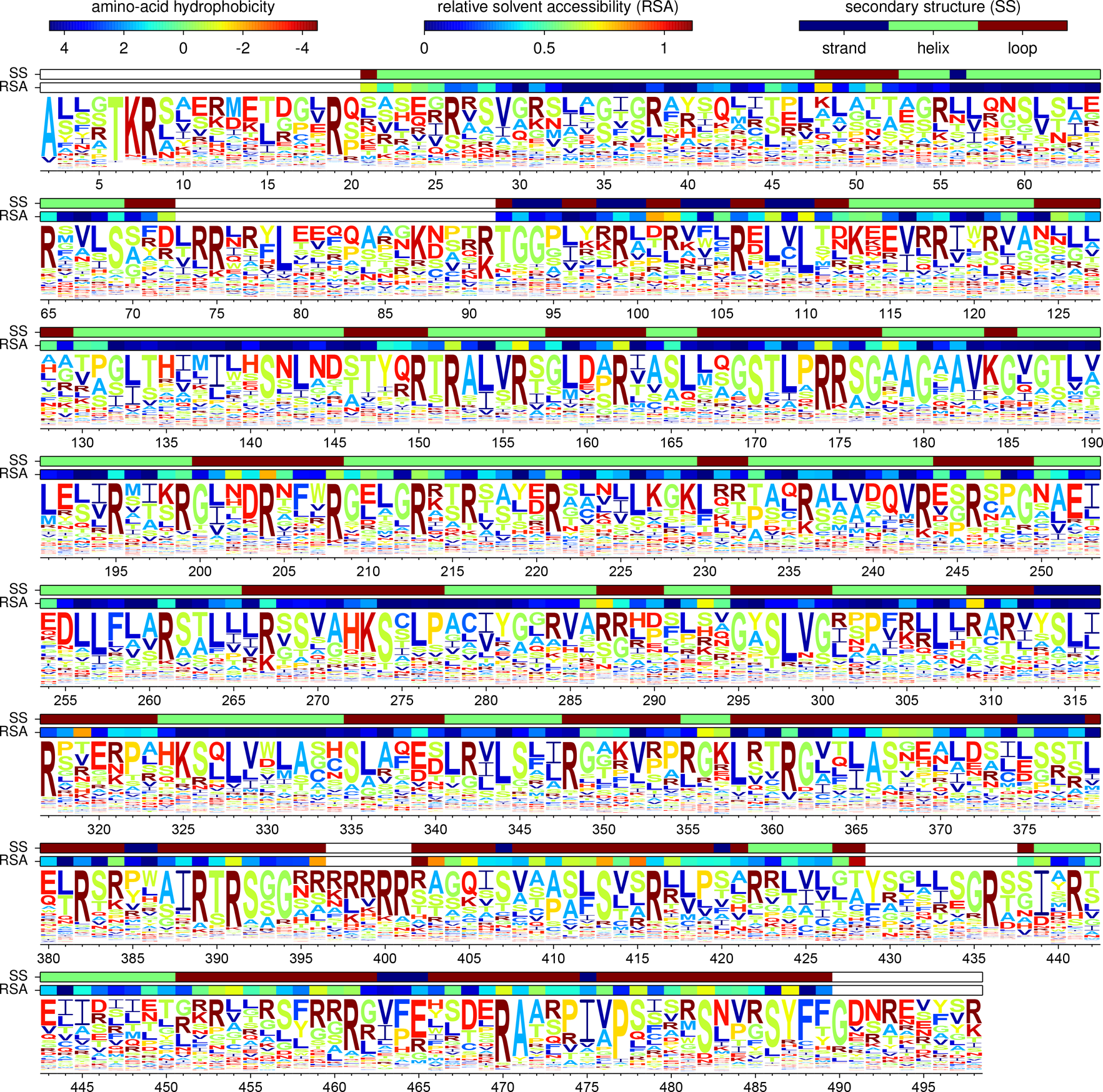
The expected frequencies of the amino acids at evolutionary equilibrium using the experimentally determined evolutionary model from passage 1 of the combined replicates and Equation 3 for the fixation probabilities. Note that these expected frequencies are slightly different than the amino-acid preferences in Figure 5D due to the structure of the genetic code. For instance, when arginine and lysine have equal preferences at a site, arginine will tend to have a higher evolutionary equilibrium frequency because it is encoded by more codons. The numerical data are in Supplementary file 2. The computer code used to generate this plot is at http://jbloom.github.io/phyloExpCM/example_2013Analysis_Influenza_NP_Human_1918_Descended.html.

### Inference of site-specific amino-acid preferences

Qualitatively, it is obvious that changes in mutation frequencies between the plasmid mutant libraries and the resulting mutant viruses reflect selection. But it is less obvious how to quantitatively analyze this information. Selection acts on the full genomes of all viruses in the population. In contrast, the experiments only measure site-independent mutation frequencies averaged over the population. Here I have analyzed this data by assuming that each site has an inherent preference for each possible amino acid. The motivation for envisioning site-heterogenous but site-independent amino-acid preferences comes from experiments suggesting that the dominant constraint on mutations that fix during NP evolution relates to protein stability (Gong et al., 2013), and that mutational effects on stability tend to be conserved in a site-independent manner (Ashenberg et al., 2013). Because the experiments generally examine each mutation in combination with several other mutations (the average clone has between 2 and 3 codon mutations; Figure 1), the site-specific amino-acid preferences are not simply selection coefficients for specific mutations. Instead, they reflect the effect of each mutation averaged over a set of genetic backgrounds.

Specifically, let *π_r,a_* denote the preference of site *r* for amino-acid *a*, with 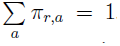. Figure 3 indicates that most observed mutations are the result of the desired codon mutagenesis, but that there is also a low rate of apparent mutations arising from Illumina sequencing errors and reverse transcription. The expected frequency *f_r,x_* of mutant codon *x* at site *r* in the mutant viruses is related to the preference 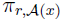 for its encoded amino-acid 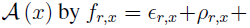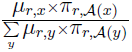 where *μ_r,x_* is the frequency that site *r* is mutagenized to codon *x* in the plasmid mutant library, *ε_r,x_* is the frequency the site is erroneously identified as *x* during sequencing, *ρ_r,x_* is the frequency the site is mutated to *x* during reverse transcription, *y* is summed over all codons, and the probability that a site experiences multiple mutations or errors in the same clone is taken to be negligibly small. The observed codon counts are multinomially distributed around these expected frequencies, so by placing a symmetric Dirichlet-distribution prior over *π_r,a_* and jointly estimating the error (*ε_r,x_* and *ρ_r,x_*) and mutation (*µ_r,x_*) rates from the appropriate samples in Figure 2, it is possible to infer the posterior mean for all amino-acid preferences by MCMC (see the Methods section).

A basic check on the consistency of the overall experimental and computational approach is to compare the amino-acid preferences inferred from different replicates, or different viral passages of the same replicate. Figure 5A,B shows that the preferences inferred from the first and second viral passages within each replicate are extremely similar, indicating that most selection occurs during initial viral creation and passage and that technical variation (preparation of samples, stochasticity in sequencing, etc) has little impact. A more crucial comparison is between the preferences inferred from the two independent experimental replicates. This comparison (Figure 5C) shows that preferences from the independent replicates are substantially but less perfectly correlated – probably the imperfect correlation is because the mutant viruses created by reverse genetics independently in each replicate are different incomplete samples of the many clones in the plasmid mutant libraries. Nonetheless, the substantial correlation between replicates shows that the sampling is sufficient to clearly reveal inherent preferences despite these experimental imperfections. Presumably better inferences can be made by aggregating data via averaging of the preferences from both replicates. Figure 5D shows such average preferences from the first passage of both replicates. These preferences are consistent with existing knowledge about NP function and stability. For example, at the conserved residues in NP's RNA binding interface (Ye et al., 2006), the amino acids found in natural sequences tend to be the ones with the highest preferences (Table 4). Similarly, for mutations that have been experimentally characterized as having large effects on NP protein stability (Gong et al., 2013; Ashenberg et al., 2013), the stabilizing amino acid has the higher preference (Table 5).

**Figure 7:**
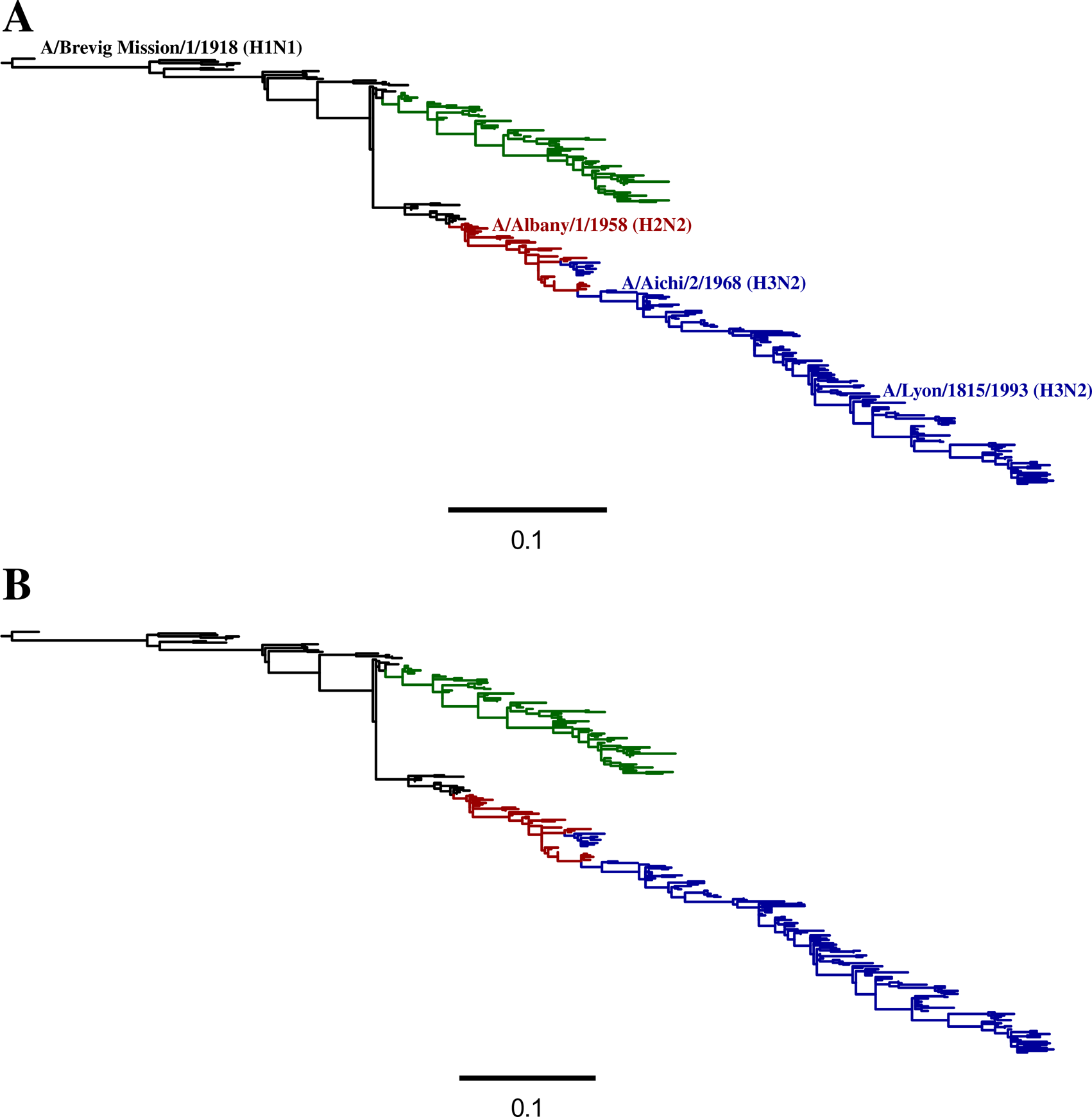
Phylogenetic tree of NPs from human influenza descended from a close relative of the 1918 virus. Black - H1N1 from 1918 lineage; green - seasonal H1N1; red - H2N2; blue - H3N2. Maximum-likelihood trees constructed using codonPhyML (Gil et al., 2013) with **(A)** the *GY94* substitution model or **(B)** the *KOSI07*+*F* substitution model. Up to three NP sequences per year from each subtype were used to build the tree. The A/Aichi/2/1968 NP that was the subject of this experiment was not one of the NP sequences randomly subsampled for the tree, so its name is indicated close to a nearly identical sequence that is shown in the tree. The computer code used to generate this tree is at http://jbloom.github.io/phyloExpCM/example_2013Analysis_Influenza_NP_Human_1918_Descended.html.

**Table 4:** For residues involved in NP's RNA-binding groove, the preferences and expected evolutionary equilibrium frequencies from the experiments correlate well with the amino-acid frequencies in naturally occurring sequences. Shown are the 17 residues in the NP RNA-binding groove in (Ye et al., 2006). The second column gives the frequencies of amino acids in all 21,108 full-length NP sequences from influenza A (excluding bat lineages) in the Influenza Virus Resource as of January-31-2014. The third column gives the experimentally measured amino-acid preferences (Figure 5D). The fourth column gives the expected evolutionary equilibrium frequency of the amino acids Figure 6). Only residues with frequencies or preferences ≥ 0.05 are listed. In all cases, the most abundant amino acid in the natural sequences has the highest expected evolutionary equilibrium frequency. In 15 of 17 cases, the most abundant amino acid in the natural sequences has the highest experimentally measured preference – in the other two cases, the most abundant amino acid in the natural sequences is among those with the highest preference.

| residue | frequencies in natural sequences | experimentally measured amino-acid preferences | expected equilibrium evolutionary frequencies from experiments |
| --- | --- | --- | --- |
| 65 | R (0.83), K (0.17) | R (0.40), K (0.10), N (0.06) | R (0.58), S (0.07) |
| 150 | R (1.00) | R (0.46), K (0.06), P (0.05), L (0.05) | R (0.63), L (0.07) |
| 152 | R (1.00) | R (0.52), K (0.07), Q (0.07) | R (0.71) |
| 156 | R (1.00) | R (0.52), Q (0.06) | R (0.69), S (0.06) |
| 174 | R (1.00) | R (0.58), N (0.06), T (0.05) | R (0.75) |
| 175 | R (1.00) | R (0.46), K (0.16) | R (0.66), K (0.08), S (0.05) |
| 195 | R (1.00) | R (0.51) | R (0.69) |
| 199 | R (1.00) | R (0.44), M (0.08), Y (0.06), V (0.05) | R (0.64), V (0.05) |
| 213 | R (1.00) | R (0.51), N (0.06) | R (0.69) |
| 214 | R (0.72), K (0.28) | K (0.24), H (0.09), R (0.09), Q (0.08), M (0.06), N (0.06), A (0.06), I (0.06) | R (0.19), K (0.17), A (0.09), H (0.07), I (0.06), L (0.06), Q (0.06) |
| 221 | R (1.00) | R (0.46), E (0.07), K (0.07) | R (0.66), L (0.05) |
| 236 | R (0.94), K (0.06) | K (0.32), R (0.30) | R (0.51), K (0.18) |
| 355 | R (1.00) | R (0.29), L (0.13), K (0.09) | R (0.43), L (0.19) |
| 357 | K (0.56), Q (0.44) | K (0.38), E (0.09), N (0.07), Y (0.05) | K (0.31), R (0.09), E (0.08), N (0.06) |
| 361 | R (1.00) | R (0.53), V (0.13) | R (0.68), V (0.11) |
| 391 | R (1.00) | R (0.59), K (0.09) | R (0.77) |
| 148 | Y (1.00) | Y (0.54), I (0.06) | Y (0.44), I (0.07), T (0.07), P (0.06), S (0.06) |

**Table 5:**
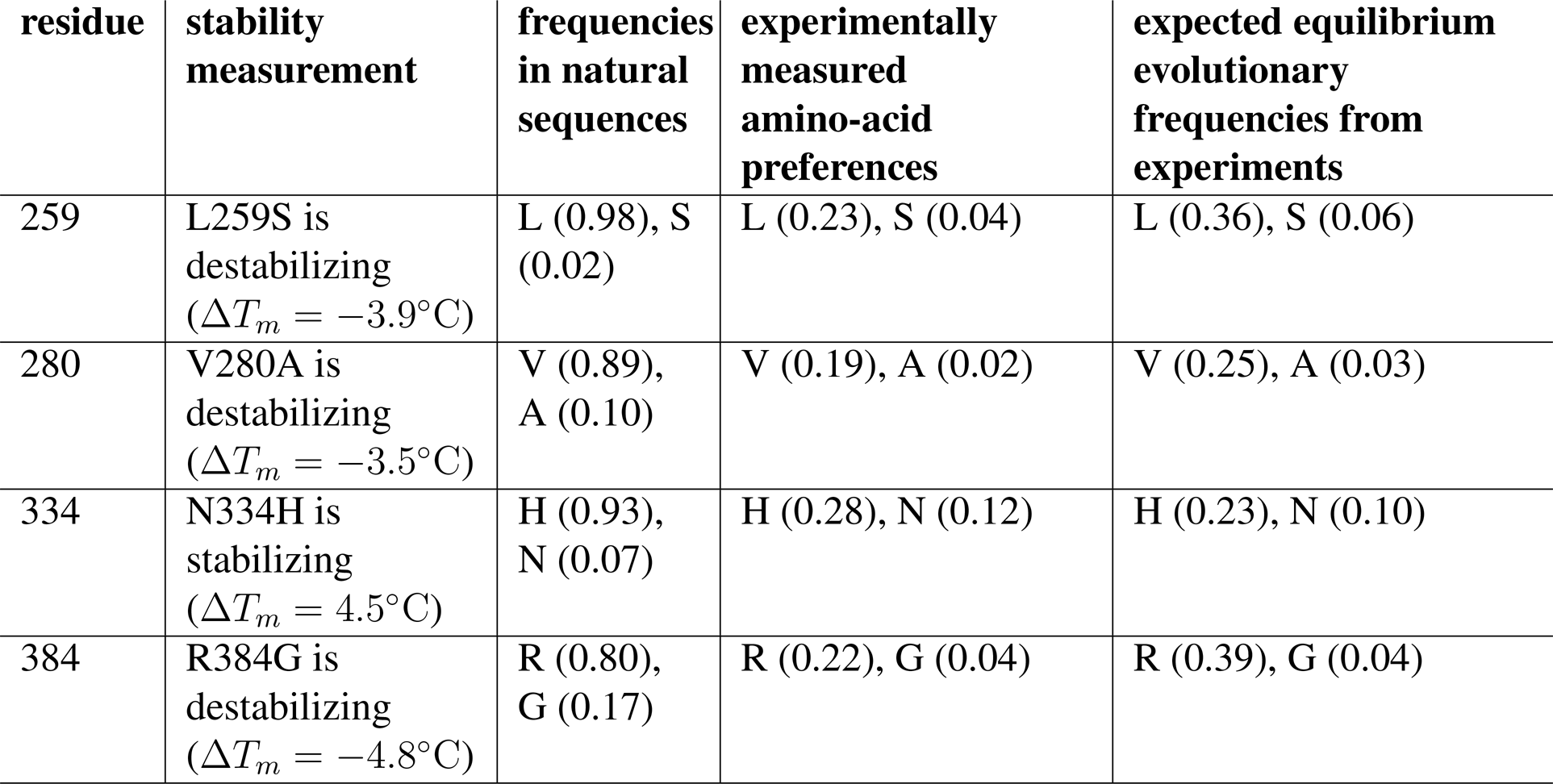
For residues where mutations have previously been characterized as having large effects on the stability of the A/Aichi/2/1968 NP, the more stable amino acid has a higher preference and is also more frequent in actual NP sequences. The second column gives the experimentally measured change in melting temperature (Δ*T_m_*) induced by the mutation to the A/Aichi/2/1968 NP as measured in (Gong et al., 2013); these mutational effects on stability are largely conserved in other NPs (Ashenberg et al., 2013). The third column gives the frequencies of the amino acids in all 21,108 full-length NP sequences from influenza A (excluding bat lineages) in the Influenza Virus Resource as of January-31-2014. The fourth column gives the experimentally measured amino-acid preferences (Figure 5D). The fifth column gives the expected evolutionary equilibrium frequency of the amino acids (Figure 6).

### The experimentally determined evolutionary model

The final step is to use the amino-acid preferences to estimate the fixation probabilities *F_r,xy_*, which can then be combined with the mutation rates to create a fully experimentally determined evolutionary model. Intuitively, it is obvious that the amino-acid preferences provide information about the fixation probabilities. For instance, it seems reasonable to expect that a mutation from *x* to *y* at site *r* will be more likely to fix (relatively larger value of *F_r,xy)_* if amino-acid 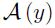 is preferred to 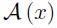 at this this site (if *π_r,A_*_(_*_y_*_)_ > *π_r,A_*_(_*_x_*_)_) and less likely to if 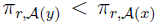. However, the exact relationship between the amino-acid preferences and the fixation probabilities is unclear. A rigorous derivation would require knowledge of unknown and probably unmeasurable population-genetics parameters for the both the deep mutational scanning experiment and the naturally evolving populations that gave rise to the sequences being analyzed phylogenetically. Instead, I provide two heuristic relationships. Both relationships satisfy detailed balance (reversibility), such that 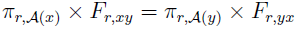, meaning that *F_r,xy_* defines a Markov process with 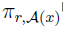 proportional to its stationary state when all amino-acid interchanges are equally probable.

It is helpful to first consider what the amino-acid preferences values actually represent. Most NP variants in the deep mutational scanning libraries contain multiple mutations, so the amino-acid preferences represent the mutational effects averaged over the nearby genetic neighborhood of the parent protein. Therefore, one interpretation is that a preference is proportional to the fraction of genetic backgrounds in which a mutation is tolerated, such that a mutation from *x* to *y* is always tolerated if 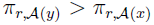, but is only sometimes tolerated if 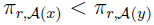. In this interpretation, there should be strong selection during initial viral growth depending on whether the mutation is tolerated in the particular genetic background in which it occurs, and then there should be little further enrichment or depletion during subsequent viral passages – loosely consistent with Figure 5A,B, which shows that the amino-acid preferences inferred after two viral passages are very similar to those inferred after one passage. Note that this interpretation can be related to the the selection-threshold evolutionary dynamics described in (Bloom et al., 2007). An equation that describes this scenario is

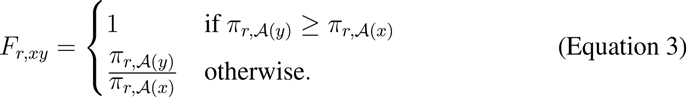

This equation is equivalent to the Metropolis acceptance criterion (Metropolis et al., 1953).

An alternative interpretation is that 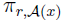 reflects the selection coefficient for the amino-acid 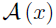 at site *r*. In this case, if the *π_r,a_* values represent the expected amino-acid equilibrium frequencies in a hypothetical evolving population in which all amino-acid interchanges are equally likely, and assuming (probably unrealistically) that this hypothetical population and the actual population in which NP evolves are in the weak-mutation limit (i.e. the population is mostly homogenous, see Desai and Fisher, 2007) and have identical constant effective population sizes, then Halpern and Bruno (1998) derive

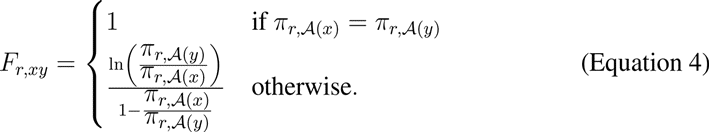

Given one of these definitions for the fixation probabilities and the mutation rates defined by Equation 2, the experimentally determined evolutionary model is defined by Equation 1. For the mutation rates and fixation probabilities used here, this evolutionary model defines a stochastic process with a unique stationary state for each site *r*. These stationary states give the expected amino-acid frequencies at evolutionary equilibrium. These evolutionary equilibrium frequencies are shown in Figure 6, and are somewhat different than the amino-acid preferences since they also depend on the structure of the genetic code (and the mutation rates when these are non-symmetric). For example, if arginine and lysine have equal preferences at a site, arginine will be more evolutionarily abundant since it has more codons.

### Phylogenetic analyses

The experimentally determined evolutionary model can be used to compute phylogenetic likelihoods, thereby enabling its comparison to existing models. In order to perform these comparisons, I first used *codonPhyML* (Gil et al., 2013) to infer maximum-likelihood trees (Figure 7) for NP sequences from human influenza using the Goldman-Yang (*GY94*) (Goldman and Yang, 1994) and the Kosiol et al. (*KOSI07*+*F*) (Kosiol et al., 2007) codon substitution models. These tree topologies were then fixed, and the branch lengths and model parameters were optimized by maximum likelihood for each of the models.

These models differ in their number of free parameters. A “free parameter” is any variable with a value that is determined from the same naturally occurring NP sequences that are being analyzed phylogenetically. The experimentally determined evolutionary model has no free parameters, since all of the properties of this model were determined by experiments that did not utilize information from naturally occurring NP sequences (the amino-acid preferences are inferred from the experiments using a symmetric prior, so in the absence of experimental data all 20 amino acids would be inferred as equally preferable at each site). Likewise, although the *KOSI07*+*F* model has a large number of exchangeability variables that were determined empirically, these variables are not free parameters since they were specified ahead of time from analysis of a general set of gene homologs that did not include NP. However, both *GY94* and *KOSI07*+*F* also contain free parameters that are estimated from the NP sequences that are being analyzed phylogenetically. In the simplest form, *GY94* contains 11 such free parameters (9 equilibrium frequencies plus transition-transversion and synonymous-nonsynonymous ratios), while *KOSI07*+*F* contains 62 parameters (60 frequencies plus transition-transversion and synonymous-nonsynonymous ratios). More complex variants add parameters allowing variation in substitution rate (Yang, 1994) or synonymous-nonsynonymous ratio among sites or lineages (Yang et al., 2000; Yang and Nielsen, 1998). For all these models, *HYPHY* (Pond et al., 2005) was used to calculate the likelihood after optimizing branch lengths and model parameters on the fixed tree topologies.

Comparison of these likelihoods strikingly validates the superiority of the experimentally determined model (Table 6 and Table 7). Adding free parameters generally improves a model's fit to data, and this is true within *GY94* and *KOSI07*+*F*. But the parameter-free experimentally determined evolutionary model describes the sequence phylogeny with a likelihood far greater than even the most highly parameterized *GY94* and *KOSI07*+*F* variants. Interpreting the amino-acid preferences as the fraction of genetic backgrounds that tolerate a mutation (Equation 3) outperforms interpreting them as selection coefficients (Equation 4), although either interpretation yields evolutionary models for NP far superior to *GY94* or *KOSI07*+*F*. Comparison using Aikake information content (AIC) to penalize parameters (Posada and Buckley, 2004) even more emphatically highlights the superiority of the experimentally determined models.

**Table 6:** Likelihoods computed using various evolutionary models after optimizing the branch lengths for the fixed tree topology inferred using the *GY94* model (Figure 7). Experimentally determined models vastly outperform *GY94* or *KOSI07*+*F*. Models are sorted by ΔAIC (Posada and Buckley, 2004), but note that the experimentally determined models all have much higher log likelihoods even before penalizing parameters. The experimentally determined models fit best if the amino-acid preferences are interpreted as the fraction of genetic backgrounds that tolerate a mutation (Equation 3) rather than as selection coefficients (Equation 4). Randomizing the experimentally determined preferences among sites makes the models far worse. All variants of *GY94* and *KOSI07*+*F* contain empirical equilibrium frequencies plus a transitiontransversion ratio and synonymous-nonsynonymous ratio (*ω*) optimized by likelihood. Some variants allow *ω* to vary across sites using discrete categories (M2a), a gamma distribution (M5), or a beta distribution plus a category (M8). Some variants allow a different *ω* for each branch. Some variants allow the rate of substitution to be gamma distributed.

| model | ΔAIC | log likelihood | parameters<br>(optimized +<br>empirical) |
| --- | --- | --- | --- |
| experimental, combined replicates | 0.0 | -12338.9 | 0 (0 + 0) |
| experimental, replicate A | 67.9 | -12372.8 | 0 (0 + 0) |
| experimental, replicate B | 106.1 | -12392.0 | 0 (0 + 0) |
| Halpern & Bruno, combined replicates | 357.9 | -12517.9 | 0 (0 + 0) |
| Halpern & Bruno, replicate A | 393.0 | -12535.4 | 0 (0 + 0) |
| Halpern & Bruno, replicate B | 455.5 | -12566.7 | 0 (0 + 0) |
| *GY94*, beta $\omega$ plus positive, one rate (M8) | 1136.8 | -12893.3 | 14 (5 + 9) |
| *GY94*, three-category $\omega$, one rate (M2a) | 1209.5 | -12929.7 | 14 (5 + 9) |
| *GY94*, gamma $\omega$, one rate (M5) | 1218.0 | -12935.9 | 12 (3 + 9) |
| *GY94*, one $\omega$, gamma rates | 1485.7 | -13069.8 | 12 (3 + 9) |
| *KOSI07+F*, three-category $\omega$, one rate (M2a) | 1679.7 | -13113.8 | 65 (5 + 60) |
| *KOSI07+F*, M8 rates-one | 1680.5 | -13114.1 | 65 (5 + 60) |
| *GY94*, one $\omega$, one rate (M0) | 1754.1 | -13205.0 | 11 (2 + 9) |
| *KOSI07+F*, gamma $\omega$, one rate | 1757.7 | -13154.8 | 63 (3 + 60) |
| *KOSI07+F*, one $\omega$, gamma rates | 1831.1 | -13191.5 | 63 (3 + 60) |
| *GY94*, branch-specific $\omega$, gamma rates (M5) | 1972.3 | -12769.1 | 556 (547 + 9) |
| *KOSI07+F*, one $\omega$, one rate (M0) | 2254.2 | -13404.0 | 62 (2 + 60) |
| *KOSI07+F*, branch-specific $\omega$, gamma rates | 2319.5 | -12891.7 | 607 (547 + 60) |
| randomized experimental, combined replicates | 3741.0 | -14209.4 | 0 (0 + 0) |
| randomized experimental, replicate A | 3809.6 | -14243.7 | 0 (0 + 0) |
| randomized experimental, replicate B | 3840.4 | -14259.1 | 0 (0 + 0) |
| randomized Halpern & Bruno, combined replicates | 4388.7 | -14533.3 | 0 (0 + 0) |
| randomized Halpern & Bruno, replicate B | 4559.1 | -14618.5 | 0 (0 + 0) |
| randomized Halpern & Bruno, replicate A | 4622.1 | -14649.9 | 0 (0 + 0) |

**Table 7:** Likelihoods for the various evolutionary models for the tree topology inferred with *codonPhyML* using *KOSI07*+*F*. This table differs from Table 6 in that it optimizes the likelihoods on the tree topology inferred with *KOSI07*+*F* rather than *GY94*.

| model | $\Delta\text{AIC}$ | log likelihood | parameters<br>(optimized +<br>empirical) |
| --- | --- | --- | --- |
| experimental, combined replicates | 0.0 | -12334.6 | 0 (0 + 0) |
| experimental, replicate A | 67.9 | -12368.5 | 0 (0 + 0) |
| experimental, replicate B | 106.2 | -12387.7 | 0 (0 + 0) |
| Halpern & Bruno, combined replicates | 356.8 | -12513.0 | 0 (0 + 0) |
| Halpern & Bruno, replicate A | 391.5 | -12530.3 | 0 (0 + 0) |
| Halpern & Bruno, replicate B | 454.8 | -12562.0 | 0 (0 + 0) |
| *GY94*, beta $\omega$ plus positive, one rate (M8) | 1183.4 | -12912.3 | 14 (5 + 9) |
| *GY94*, three-category $\omega$, one rate (M2a) | 1209.4 | -12925.3 | 14 (5 + 9) |
| *GY94*, gamma $\omega$, one rate (M5) | 1219.6 | -12932.4 | 12 (3 + 9) |
| *GY94*, one $\omega$, gamma rates | 1493.1 | -13069.1 | 12 (3 + 9) |
| *KOSI07+F*, three-category $\omega$, one rate (M2a) | 1676.0 | -13107.6 | 65 (5 + 60) |
| *KOSI07+F*, M8 rates-one | 1676.6 | -13107.9 | 65 (5 + 60) |
| *KOSI07+F*, gamma $\omega$, one rate | 1753.3 | -13148.2 | 63 (3 + 60) |
| *GY94*, one $\omega$, one rate (M0) | 1762.2 | -13204.7 | 11 (2 + 9) |
| *KOSI07+F*, one $\omega$, gamma rates | 1834.3 | -13188.7 | 63 (3 + 60) |
| *GY94*, branch-specific $\omega$, gamma rates (M5) | 1980.8 | -12769.0 | 556 (547 + 9) |
| *KOSI07+F*, one $\omega$, one rate (M0) | 2256.8 | -13401.0 | 62 (2 + 60) |
| *KOSI07+F*, branch-specific $\omega$, gamma rates | 2324.0 | -12889.6 | 607 (547 + 60) |
| randomized experimental, combined replicates | 3741.3 | -14205.2 | 0 (0 + 0) |
| randomized experimental, replicate A | 3809.4 | -14239.3 | 0 (0 + 0) |
| randomized experimental, replicate B | 3841.4 | -14255.3 | 0 (0 + 0) |
| randomized Halpern & Bruno, combined replicates | 4387.6 | -14528.4 | 0 (0 + 0) |
| randomized Halpern & Bruno, replicate B | 4557.9 | -14613.6 | 0 (0 + 0) |
| randomized Halpern & Bruno, replicate A | 4620.8 | -14645.0 | 0 (0 + 0) |

There is also a clear correlation between the quality and volume of experimental data and the phylogenetic fit: models from individual experimental replicates give lower likelihoods than both replicates combined, and the technically superior *replicate A* (recall the comparison in Figure 4) gives a better likelihood than *replicate B* (Table 6 and Table 7). This fact suggests that improvements in experimental methodology that improve the accuracy of the measured mutational effects should lead to even better experimentally determined evolutionary models.

In Table 6 and Table 7, the site-specific experimentally determined model is compared to variants of two general models (*GY94* and *KOSI07*+*F*) that apply broadly to all proteins. More recently, it has become possible to estimate non-site-specific (identical across sites) codon and amino-acid models using naturally occurring sequences from specific proteins or viruses (Dang et al., 2010; De Maio et al., 2013). One could therefore ask if the experimentally determined model is superior because it is site-specific, or simply because it is experimentally derived from deep mutational scanning of influenza. To address this question, I created “randomized” experimentally determined models in which the deep mutational scanning data was randomly shuffled among protein sites. These randomized models are still derived from deep mutational scanning of influenza, but have lost their linkage to site-specific experimental information. These randomized models are greatly inferior to all of the other models considered here (Table 6, Table 7). Therefore, the superiority of the experimentally determined model is due to its utilization of site-specific information from the deep mutational scanning – if this site specificity is lost, the model becomes far worse than general models such as GY94 or *KOSI07*+*F*.

## Discussion

These results establish that an experimentally determined evolutionary model is far superior to existing models for describing the phylogeny of NP gene sequences. The extent of this superiority is striking. The parameter-free evolutionary model dramatically outperforms even the most highly parameterized existing models using the parameter-penalizing metric of AIC – but more remarkably, it also outperforms these parameterized models by over 400 log-likelihood units even in the absence of parameter penalization (Table 6). The reason for this superiority is easy to understand: proteins have strong and fairly conserved preferences for specific amino acids at different sites (Ashenberg et al., 2013), but these site-specific preferences are ignored by most existing phylogenetic models. Inspection of the overlaid bars in Figure 5D illustrates the inadequacy of trying to capture these preferences simply by classifying sites based on gross features of protein structure (Thorne et al., 1996; Goldman et al., 1998) – the site-specific amino-acid preferences are not simply related to secondary structure or solvent accessibility. The complexity of the preferences in Figure 5D also show the limitations of attempting to infer aminoacid preference parameters for a smal number of site classes from sequence data (Lartillot and Philippe, 2004; Le et al., 2008; Wu et al., 2013; Wang et al., 2008), as it is clear that each site is unique. Direct experimental measurement therefore represents a highly attractive method for determining the idiosyncratic constraints that affect the evolution of each site in a gene.

Another appealing aspect of an experimentally determined evolutionary model is interpretability. A frustrating aspect of existing evolutionary models is the inability to interpret many of their free parameters directly in evolutionary or molecular terms. For example, the equilibrium frequency parameters used by most existing models reflect some unknown combination of mutational bias and selection for specific codons or amino acids – but the relative contributions of these factors in determining the parameter values is unclear. On the other hand, all aspects of the experimentally determined evolutionary model can be related to direct measurements, making them more amenable to direct interpretation. So even if such a model were eventually augmented with a few free parameters, this could be done in a way that allowed these parameters to retain a clear connection to the molecular processes of biology and evolution.

The results presented here also demonstrate that phylogenetic evolutionary models can be greatly improved while retaining the assumption of independence of sites. Phylogenetic evolutionary models make two assumptions that are egregiously bad from the perspective of the protein chemist: first these models assume that sites are identical (or at least can be described by a small number of classes), and second they assume that sites are independent. The experimentally determined model eliminates the first assumption, but does nothing to relax the second. Is this model therefore inconsistent with the idea that epistasis is common during protein evolution (Lunzer et al., 2010)? In fact, experiments show that a general conservation of site-specific amino-acid preferences is entirely consistent with epistasis. For instance, there is known epistasis among some of the mutations fixed along the NP phylogenetic tree analyzed here (Gong et al., 2013) – but the site-specific compatibilities of amino acids with the protein’s structural stability are largely conserved among homologs on this tree, even for sites involved in epistatic interactions (Ashenberg et al., 2013). The reason is that evolutionary relevant epistasis can arise from subtle and transient fluctuations in properties such as protein stability, whereas the phylogenetic improvements from a site-specific model probably come mostly from capturing basic information about the compatibility of amino acids with a protein’s evolutionarily conserved structure. Models that assume independence among sites can therefore still lead to major improvements if the site-specific amino-acid preferences are accurately represented.

The major drawback of the experimentally determined evolutionary model is its lack of generality. While this model is clearly superior for influenza NP, it is entirely unsuitable for other genes. At first blush, it might seem that the arduous experiments described here provide data that is unlikely to ever become available for most situations of interest. However, it is worth remembering that today’s arduous experiment frequently becomes routine in a few years. For example, the very gene sequences that are the subjects of molecular phylogenetics were once rare pieces of data – now such sequences are so abundant that they easily overwhelm modern computers. The experimental ease of the deep mutational scanning approach used here is on a comparable trajectory: similar approaches have already been applied to several proteins (Fowler et al., 2010; Melamed et al., 2013; Traxlmayr et al., 2012; Starita et al., 2013; Roscoe et al., 2013), and there continue to be rapid improvements in techniques for mutagenesis (Firnberg and Ostermeier, 2012; Jain and Varadarajan, 2014) and sequencing (Hiatt et al., 2010; Schmitt et al., 2012; Lou et al., 2013). Given these prospects for technical improvements in deep mutational scanning, it is therefore especially encouraging that the phylogenetic fit of the NP evolutionary model improves with the quality and volume of experimental data from which it is derived (Table 6). The increasing availability of similar high-throughput data for a vast range of proteins has the potential to transform phylogenetic analyses by greatly increasing the accuracy of evolutionary models, while at the same time replacing a plethora of free parameters with experimentally measured quantities that can be given clear biological and evolutionary interpretations.

## Methods

### Availability of data and computer code

Illumina sequencing data are available at the SRA (accession SRP036064, http://www.ncbi.nlm.nih.gov/sra/?term=SRP036064). A description and links to the source code used to analyze the sequencing data and infer the amino-acid preferences is at http://jbloom.github.io/mapmuts/example_2013Analysis_Influenza_NP_Aichi68.html. A description and links to the source code used for the phylogenetic analyses is at http://jbloom.github.io/phyloExpCM/example_2013Analysis_Influenza_NP_Human_1918_Descended.html.

### Experimental measurement of mutation rates

To measure mutation rates, I generated GFP-carrying viruses with all genes derived from A/WSN/1933 (H1N1) by reverse genetics as described previously (Bloom et al., 2010). These viruses were repeatedly passaged at limiting dilution in MDCK-SIAT1-CMV-PB1 cells (Bloom et al., 2010) using *low serum media* (Opti-MEM I with 0.5% heat-inactivated fetal bovine serum, 0.3% BSA, 100 U/ml penicillin, 100 *µ*g/ml streptomycin, and 100 *µ*g/ml calcium chloride) – a moderate serum concentration was retained and no trypsin was added because viruses with the WSN HA and NA are trypsin independent (Goto and Kawaoka, 1998). These passages were performed for 27 replicate populations. For each passage, 100 *µ*l containing the equivalent of 2 *µ*l of virus collection was added to the first row of a 96-well plate. The virus was serially diluted 1:5 down the plate such that at the conclusion of the dilutions, each well contained 80 *µ*l of virus dilution. MDCK-SIAT1-CMV-PB1 cells were then added to each well in a 50 *μ*l volume containing 2.5 × 10^3^ cells. The plates were grown for approximately 80 hours, and wells were examined for cytopathic effect indicative of viral growth. The last well with cytopathic effect was collected and used as the parent population for the next round of limiting-dilution passage.

After 25 limiting-dilution passages, 10 of the 27 viral populations no longer caused any visible GFP expression in the cells in which they caused cytopathic effect, indicating fixation of a mutation that ablated GFP fluorescence. The 17 remaining populations all caused fluorescence in infected cells, although in some cases the intensity was visibly reduced – these populations therefore must have retained at least a partially functional GFP. Total RNA was purified from each viral population, the PB1 segment was reverse-transcribed using the primers CATGATCGTCTCGTATTAGTAGAAACAAGGCATTTTTTCATGAAGGACAAGC and CATGATCGTCTCAGGGAGCGAAAGCAGGCAAACCATTTGATTGG, and the reverse-transcribed cDNA was amplified by conventional PCR using the same primers. For 22 of the 27 replicate viral populations, this process amplified an insert with the length expected for the full GFP-carrying PB1 segment. For 2 of the replicates, this amplified inserts between 0.4 and 0.5 kb shorter than the expected length, suggesting an internal deletion in part of the segment. For 3 replicates, this failed to amplify any insert, suggesting total loss
of the GFP-carrying PB1 segment, a very large internal deletion, or rearrangement that rendered the reverse-transcription primers ineffective. For the 24 replicates from which an insert could be amplified, the GFP coding region was Sanger sequenced to determine the consensus sequence. The results are in Table 1 and Table 2.

To estimate *R_m→n_*, it is necessary to normalize by the nucleotide composition of the GFP gene. The numbers of each nucleotide in this gene are: *N_A_* = 175, *N_T_* = 103, *N_C_* = 241, and *N_G_* = 201. Given that the counts in Table 2 come after 25 passages of 24 replicates:

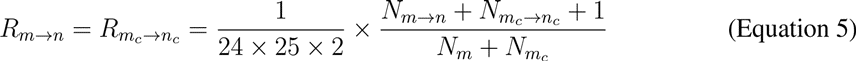

where *N*_*m → n*_ is the number of observed mutations from *m* to *n* in Table 2, m_c_ indicates the complement of DNA nucleotide *m* (so for example *A_c_* = *T*). The one in the numerator is a pseudocount added to the observed counts of each type of mutation in order to avoid estimating rates of zero. The values of *R*_*m → n*_ estimated from Equation 5 give the probability that a nucleotide that is already *m* will mutate to *n* in a single tissue-culture generation.

### Construction of NP codon-mutant libraries

The goal was to construct a mutant library with an average of two to three random codon mutations per gene. Most techniques for creating mutant libraries of full-length genes, such as error-prone PCR (Cirino et al., 2003) and chemical mutagenesis (Neylon, 2004), introduce mutations at the nucleotide level, meaning that codon substitutions involving multiple nucleotide changes occur at a negligible rate. Recently, several groups have developed strategies for introducing codon mutations along the lengths of entire genes (Firnberg and Ostermeier, 2012; Jain and Varadarajan, 2014, J. Kitzman and J. Shendure - personal communication). Most of these strategies are designed to create exactly one codon mutation per gene. For my experiments, it was desirable to introduce a distribution of around one to four codon mutations per gene to examine the average effects of mutations in a variety of closely related genetic backgrounds. Therefore, I devised a codon-mutagenesis protocol specifically for this purpose.

This technique involved iterative rounds of low-cycle PCR with pools of mutagenic synthetic oligonucleotides that each contain a randomized NNN triplet at a specific codon site. Two replicate libraries each of the wildtype and N334H variants of the Aichi/1968 NP were prepared in full biological duplicate, beginning each with independent preps of the plasmid templates pHWAichi68-NP and pHWAichi68-NP-N334H. The sequences of the NP genes in these plasmids are provided in (Gong et al., 2013). To avoid cross-contamination, all purification steps used an independent gel for each sample, with the relevant equipment thoroughly washed to remove residual DNA.

First, for each codon except for that encoding the initiating methionine in the 498-residue NP gene, I designed an oligonucleotide that contained a randomized NNN nucleotide triplet preceded by the 16 nucleotides upstream of that codon in the NP gene and followed by the 16 nucleotides downstream of that codon in the NP gene. I ordered these oligonucleotides in 96-well plate format from Integrated DNA Technologies, and combined them in equimolar quantities to create the *forward-mutagenesis* primer pool. I also designed and ordered the reverse complement of each of these oligonucleotides, and combined them in equimolar quantities to create the *reverse-mutagenesis* pool. The primers for the N334H variants differed only for those that overlapped the N334H codon. I also designed end primers that annealed to the termini of the NP sequence and contained sites appropriate for BsmBI cloning into the influenza reverse-genetics plasmid pHW2000 (Hoffmann et al., 2000). These primers are *5’-BsmBI-Aichi68-NP* (catgatcgtctcagggagcaaaagcagggtagataatcactcacag) and *3’-BsmBI-Aichi68-NP* (catgatcgtctcgtattagtagaaacaagggtatttttcttta).

I set up PCR reactions that contained 1 *µ*l of 10 ng/*µ*l template pHWAichi68-NP plasmid (Gong et al., 2013), 25 *µ*l of 2X KOD Hot Start Master Mix (product number 71842, EMD Millipore), 1.5 *µ*l each of 10 *µ*M solutions of the end primers *5’-BsmBI-Aichi68-NP* and *3’- BsmBI-Aichi68-NP*, and 21 *µ*l of water. I used the following PCR program (referred to as the *amplicon PCR program* in the remainder of this paper):

1. 95°C for 2 minutes
2. 95°C for 20 seconds
3. 70°C for 1 second
4. 50°C for 30 seconds cooling to 50°C at 0.5°C per second.
5. 70°C for 40 seconds
6. Repeat steps 2 through 5 for 24 additional cycles
7. Hold 4°C

The PCR products were purified over agarose gels using ZymoClean columns (product number D4002, Zymo Research) and used as templates for the initial codon mutagenesis fragment PCR.

Two fragment PCR reactions were run for each template. The forward-fragment reactions contained 15 *µ*l of 2X KOD Hot Start Master Mix, 2 *µ*l of the *forward mutagenesis* primer pool at a total oligonucleotide concentration of 4.5 *µ*M, 2 *µ*l of 4.5 *µ*M *3’-BsmBI-Aichi68-NP*, 4 *µ*l of 3 ng/*µ*l of the aforementioned gel-purified linear PCR product template, and 7 *µ*l of water. The reverse-fragment reactions were identical except that the *reverse mutagenesis* pool was substituted for the *forward mutagenesis* pool, and that *5’-BsmBI-Aichi68-NP* was substituted for *3’-BsmBI-Aichi68-NP*. The PCR program for these fragment reactions was identical to the *amplicon PCR program* except that it utilized a total of 7 rather than 25 thermal cycles.

The products from the fragment PCR reactions were diluted 1:4 in water. These dilutions were then used for the joining PCR reactions, which contained 15 *µ*l of 2X KOD Hot Start Master Mix, 4 *µ*l of the 1:4 dilution of the forward-fragment reaction, 4 *µ*l of the 1:4 dilution of the reverse-fragment reaction, 2 *µ*l of 4.5 *µ*M *5’-BsmBI-Aichi68-NP*, 2 *µ*l of 4.5 *µ*M *3’-BsmBI- Aichi68-NP*, and 3 *µ*l of water. The PCR program for these joining reactions was identical to the *amplicon PCR program* except that it utilized a total of 20 rather than 25 thermal cycles. The products from these joining PCRs were purified over agarose gels.

The purified products of the first joining PCR reactions were used as templates for a second round of fragment reactions followed by joining PCRs. These second-round products were used as templates for a third round. The third-round products were purified over agarose gels, digested with BsmBI (product number R0580L, New England Biolabs), and ligated into a dephosphorylated (Antarctic Phosphatase, product number M0289L, New England Biolabs) BsmBI digest of pHW2000 (Hoffmann et al., 2000) using T4 DNA ligase. The ligations were purified using ZymoClean columns, electroporated into ElectroMAX DH10B T1 phage-resistant competent cells (product number 12033-015, Invitrogen) and plated on LB plates supplemented with 100 *µ*g/ml of ampicillin. These transformations yielded between 400,000 and 800,000 unique transformants per plate, as judged by plating a 1:4,000 dilution of the transformations on a second set of plates. Transformation of a parallel no-insert control ligation yielded approximately 50-fold fewer colonies, indicating that self ligation of pHW2000 only accounts for a small fraction of the transformants. For each library, I performed three transformations, grew the plates overnight, and then scraped the colonies into liquid LB supplemented with ampicillin and mini-prepped several hours later to yield the plasmid mutant libraries. These libraries each contained in excess of 10^6^ unique transformants, most of which will be unique codon mutants of the NP gene.

I sequenced the NP gene for 30 individual clones drawn from the four mutant libraries. As shown in Figure 1, the number of mutations per clone was approximately Poisson distributed and the mutations occurred uniformly along the primary sequence. If all codon mutations are made with equal probability, 9/63 of the mutations should be single-nucleotide changes, 27/63 should be two-nucleotide changes, and 27/63 should be three-nucleotide changes. This is approximately what was observed in the Sanger-sequenced clones. The nucleotide composition of the mutated codons was roughly uniform, and there was no tendency for clustering of multiple mutations in primary sequence. The results of this Sanger sequencing are compatible with the mutation frequencies obtained from deep sequencing the **mutDNA** samples after subtracting off the sequencing error rate estimated from the **DNA** samples (Figure 3), especially considering that the statistics from the Sanger sequencing are subject to sampling error due to the limited number of clones analyzed.

### Viral growth and passage

Two independent replicates of viral growth and passage were performed (*replicate A* and *replicate B*). The procedures were similar between replicates, but there were a few small differences. In the actual experimental chronology, *replicate B* was performed first, and the modifications in *replicate A* were designed to improve the sampling of the mutations by the created mutant viruses. These modifications may be the reason why *replicate A* slightly outperforms *replicate B* by two objective measures: the viruses more completely sample the codon mutations (Figure 4), and the evolutionary model derived solely from *replicate A* gives a higher likelihood than the evolutionary model derived solely from *replicate B* (Table 6 and Table 7).

For *replicate B*, I used reverse genetics to rescue viruses carrying the Aichi/1968 NP or one of its derivatives, PB2 and PA from the A/Nanchang/933/1995 (H3N2), a PB1 gene segment encoding GFP, and HA / NA / M / NS from A/WSN/1933 (H1N1) strain. With the exception of the variants of NP used, these viruses are identical to those described in (Gong et al., 2013), and were rescued by reverse genetics in 293T-CMV-Nan95-PB1 and MDCK-SIAT1-CMV-Nan95-PB1 cells as described in that reference. The previous section describes four NP codon-mutant libraries, two of the wildtype Aichi/1968 gene (WT-1 and WT-2) and two of the N334H variant (N334H-1 and N334H-2). I grew mutant viruses from all four mutant libraries, and four paired unmutated viruses from independent preps of the parent plasmids. A major goal was to maintain diversity during viral creation by reverse genetics – the experiment would obviously be undermined if most of the rescued viruses derived from a small number of transfected plasmids. I therefore performed the reverse genetics in 15 cm tissue-culture dishes to maximize the number of transfected cells. Specifically, 15 cm dishes were seeded with 10^7^ 293T-CMV-Nan95-PB1 cells in D10 media (DMEM with 10% heat-inactivated fetal bovine serum, 2 mM L-glutamine, 100 U/ml penicillin, and 100 *µ*g/ml streptomycin). At 20 hours post-seeding, the dishes were transfected with 2.8 *µ*g of each of the eight reverse-genetics plasmids. At 20 hours post-transfection, about 20% of the cells expressed GFP (indicating transcription by the viral polymerase of the GFP encoded by pHH-PB1flank-eGFP), suggesting that many unique cells were transfected. At 20 hours post-transfection, the media was changed to the *low serum media* described above. At 78 hours post-transfection, the viral supernatants were collected, clarified by centrifugation at 2000 × g for 5 minutes, and stored at 4°C. The viruses were titered by flow cytometry as described previously (Gong et al., 2013). A control lacking the NP gene yielded no infectious virus as expected.

The virus was then passaged in MDCK-SIAT1-CMV-Nan95-PB1 cells. These cells were seeded into 15 cm dishes, and when they had reached a density of 10^7^ per plate they were infected with 10^6^ infectious particles (MOI of 0.1) of the transfectant viruses in *low serum media*. After 18 hours, 30-50% of the cells were green as judged by microscopy, indicating viral spread. At 40 hours post-transfection, 100% of the cells were green and many showed clear signs of cytopathic effect. At this time the viral supernatants were again collected, clarified, and stored at 4°C. NP cDNA isolated from these viruses was the source the deep-sequencing samples **virus-p1** and **mutvirus-p1** in Figure 2. The virus was then passaged a second time exactly as before (again using an MOI of 0.1). NP cDNA from these twice-passaged viruses constituted the source for the samples **virus-p2** and **mutvirus-p2** in Figure 2.

For *replicate A*, all viruses (both the four mutant viruses and the paired unmutated controls) were re-grown independently from the same plasmids preps used for *replicate B*. The experimental process was identical to that used for *replicate B* except for the following: Standard influenza viruses (rather than the GFP-carrying variants) were used, so plasmid pHW-Nan95-PB1 (Gong et al., 2013) was substituted for pHH-PB1flank-eGFP during reverse genetics, and 293 T and MDCK-SIAT1 cells were substituted for the PB1-expressing variants. Rather than creating the viruses by transfecting a single 15-cm dish, each sample was created by transfecting two 12-well dishes, with the dishes seeded at 3 × 10^5^ 293 T and 5 × 10^4^ MDCK-SIAT1 cells prior to transfection. The passaging was then done in four 10 cm dishes for each sample, with the dishes seeded at 4 ×10^6^ MDCK-SIAT1 cells 12-14 hours prior to infection. The passaging was still done at an MOI of 0.1. These modifications were designed to increase diversity in the viral population. These viruses were titered by TCID50 rather than flow cytometry.

### Sample preparation and Illumina sequencing

For each sample, a PCR amplicon was created to serve as the template for Illumina sequencing. The steps used to generate the PCR amplicon for each of the seven sample types (Figure 2) are listed below. Once the PCR template was generated, for all samples the PCR amplicon was created using the *amplicon PCR program* described above in 50 *µ*l reactions consisting of 25 *µ*l of 2X KOD Hot Start Master Mix, 1.5 *µ*l each of 10 *µ*M of *5’-BsmBI-Aichi68-NP* and *3’-BsmBI-Aichi68-NP*, the indicated template, and ultrapure water. A small amount of each PCR reaction was run on an analytical agarose gel to confirm the desired band. The remainder was then run on its own agarose gel without any ladder (to avoid contamination) after carefully cleaning the gel rig and all related equipment. The amplicons were excised from the gels, purified over ZymoClean columns, and analyzed using a NanoDrop to ensure that the absorbance at 260 nm was at least 1.8 times that at 230 nm and 280 nm. The templates were as follows:

- **DNA**: The templates for these amplicons were 10 ng of the unmutated independent plasmid preps used to create the codon mutant libraries.
- **mutDNA**: The templates for these amplicons were 10 ng of the plasmid mutant libraries.
- **RNA**: This amplicon quantifies the net error rate of transcription and reverse transcription. Because the viral RNA is initially transcribed from the reverse-genetics plasmids by RNA polymerase I but the bidirectional reverse-genetics plasmids direct transcription of RNA by both RNA polymerases I and II (Hoffmann et al., 2000), the RNA templates for these amplicons were transcribed from plasmids derived from pHH21 (Neumann et al., 1999), which only directs transcription by RNA polymerase I. The unmutated WT and N334H NP genes were cloned into this plasmid to create pHH-Aichi68-NP and pHH-Aichi68-NP-N334H. Independent preparations of these plasmids were transfected into 293 T cells, transfecting 2 *µ*g of plasmid into 5 × 10^5^ cells in 6-well dishes. After 32 hours, total RNA was isolated using Qiagen RNeasy columns and treated with the Ambion TURBO DNAfree kit (Applied Biosystems AM1907) to remove residual plasmid DNA. This RNA was used as a template for reverse transcription with AccuScript (Agilent 200820) using the primers *5’-BsmBI-Aichi68-NP* and *3’-BsmBI-Aichi68-NP*. The resulting cDNA was quantified by qPCR specific for NP (see below), which showed high levels of NP cDNA in the reverse-transcription reactions but undetectable levels in control reactions lacking the reverse transcriptase, indicating that residual plasmid DNA had been successfully removed. A volume of cDNA that contained at least 2 × 10^6^ NP cDNA molecules (as quantified by qPCR) was used as template for the amplicon PCR reaction. Control PCR reactions using equivalent volumes of template from the no reverse-transcriptase control reactions yielded no product.
- **virus-p1**: This amplicon was derived from virus created from the unmutated plasmid and collected at the end of the first passage. Clarified virus supernatant was ultracentrifuged at 64,000 × g for 1.5 hours at 4 °C, and the supernatant was decanted. Total RNA was then isolated from the viral pellet using a Qiagen RNeasy kit. This RNA was used as a template for reverse transcription with AccuScript using the primers *5’-BsmBI-Aichi68-NP* and *3’-BsmBI-Aichi68-NP*. The resulting cDNA was quantified by qPCR, which showed high levels of NP cDNA in the reverse-transcription reactions but undetectable levels in control reactions lacking the reverse transcriptase. A volume of cDNA that contained at least 10^7^ NP cDNA molecules (as quantified by qPCR) was used as template for the amplicon PCR reaction. Control PCR reactions using equivalent volumes of template from the no reverse-transcriptase control reactions yielded no product.
- **virus-p2, mutvirus-p1, mutvirus-p2**: These amplicons were created as for the **virus-p1** amplicons, but used the appropriate virus as the initial template as outlined in Figure 2.

An important note: it was found that the use of relatively new RNeasy kits with *β*-mercaptoethanol (a reducing agent) freshly added per the manufacturer's instructions was necessary to avoid what appeared to be oxidative damage to purified RNA.

The overall experiment only makes sense if the sequenced NP genes derive from a large diversity of initial template molecules. Therefore, qPCR was used to quantify the molecules produced by reverse transcription to ensure that a sufficiently large number were used as PCR templates to create the amplicons. The qPCR primers were *5’-Aichi68-NP-for* (gcaacagctggtctgactcaca) and *3’-Aichi68-NP-rev* (tccatgccggtgcgaacaag). The qPCR reactions were performed using the SYBR Green PCR Master Mix (Applied Biosystems 4309155) following the manufacturer’s instructions. Linear NP PCR-ed from the pHWAichi68-NP plasmid was used as a quantification standard – the use of a linear standard is important, since amplification efficiencies differ for linear and circular templates (Hou et al., 2010). The standard curves were linear with respect to the amount of NP standard over the range from 10^2^ to 10^9^ NP molecules. These standard curves were used to determine the absolute number of NP cDNA molecules after reverse transcription. Note that the use of only 25 thermal cycles in the *amplicon PCR program* provides a second check that there are a substantial number of template molecules, as this moderate number of thermal cycles will not lead to sufficient product if there are only a few template molecules.

In order to allow the Illumina sequencing inserts to be read in both directions by paired-end 50 nt reads (Supplementary figure 1), it was necessary to us an Illumina library-prep protocol that created NP inserts that were roughly 50 nt in length. This was done via a modification of the Illumina Nextera protocol. First, concentrations of the PCR amplicons were determined using PicoGreen (Invitrogen P7859). These amplicons were used as input to the Illumina Nextera DNA Sample Preparation kit (Illumina FC-121-1031). The manufacturer's protocol for the tagmentation step was modified to use 5-fold less input DNA (10 ng rather than 50 ng) and two-fold more tagmentation enzyme (10 *µ*l rather than 5 *µ*l), and the incubation at 55 °C was doubled from 5 minutes to 10 minutes. Samples were barcoded using the Nextera Index Kit for 96-indices (Illumina FC-121-1012). For index 1, the barcoding was: **DNA** with N701, **RNA** with N702, **mutDNA** with N703, **virus-p1** with N704, **mutvirus-p1** with N705, **virus-p2** with N706, and **mutvirus-p2** with N707. After completion of the Nextera PCR, the samples were subjected to a ZymoClean purification rather than the bead cleanup step specified in the Nextera protocol. The size distribution of these purified PCR products was analyzed using an Agilent 200 TapeStation Instrument. If the NP sequencing insert is exactly 50 nt in size, then the product of the Nextera PCR should be 186 nt in length after accounting for the addition of the Nextera adaptors. The actual size distribution was peaked close to this value. The ZymoClean-purified PCR products were quantified using PicoGreen and combined in equal amounts into pools: a WT-1 pool of the seven samples for that library, a WT-2 pool of the seven samples for that library, etc. These pools were subjected to further size selection by running them on a 4% agarose gel versus a custom ladder containing 171 and 196 nt bands created by PCR from a GFP template using the forward primer gcacggggccgtcgccg and the reverse primers tggggcacaagctggagtacaac (for the 171 nt band) and gacttcaaggaggacggcaacatcc (for the 196 nt band). The gel slice for the sample pools corresponding to sizes between 171 and 196 nt was excised, and purified using a ZymoClean column. A separate clean gel was run for each pool to avoid cross-contamination.

Library QC and cluster optimization were performed using Agilent Technologies qPCR NGS Library Quantification Kit (Agilent Technologies, Santa Clara, CA, USA). Libraries were introduced onto the flow cell using an Illumina cBot (Illumina, Inc., San Diego, CA, USA) and a TruSeq Rapid Duo cBot Sample Loading Kit. Cluster generation and deep sequencing was performed on an Illumina HiSeq 2500 using an Illumina TruSeq Rapid PE Cluster Kit and TruSeq Rapid SBS Kit. A paired-end, 50 nt read-length (PE50) sequencing strategy was performed in rapid run mode. Image analysis and base calling were performed using Illumina's Real Time Analysis v1.17.20.0 software, followed by demultiplexing of indexed reads and generation of FASTQ files, using Illumina's CASAVA v1.8.2 software (http://www.illumina.com/software.ilmn). These FASTQ files were uploaded to the SRA under accession SRP036064 (see http://www.ncbi.nlm.nih.gov/sra/?term=SRP036064).

### Read alignment and quantification of mutation frequencies

A custom Python software package, *mapmuts*, was created to quantify the frequencies of mutations from the Illumina sequencing. A description of the software as utilized in the present work is at http://jbloom.github.io/mapmuts/example_2013Analysis_Influenza_NP_Aichi68.html. Briefly:

1. Reads were discarded if either read in a pair failed the Illumina chastity filter, had a mean Q-score less than 25, or had more than two ambiguous (N) nucleotides.
2. The remaining paired reads were aligned to each other, and retained only if they shared at least 30 nt of overlap, disagreed at no more than one site, and matched the expected terminal Illumina adaptors with no more than one mismatch.
3. The overlap of the paired reads was aligned to NP, disallowing alignments with gaps or more than six nucleotide mismatches. A small fraction of alignments corresponded exclusively to the noncoding termini of the viral RNA; the rest contained portions of the NP coding sequence.
4. For every paired read that aligned with NP, the codon identity was called if both reads concurred for all three nucleotides in the codon. If the reads disagreed or contained an ambiguity in that codon, the identity was not called.

### Inference of the amino-acid preferences

The approach described here is based on the assumption that there is an inherent preference for each amino acid at each site in the protein. This assumption is clearly not completely accurate, as the effect of a mutation at one site can be influenced by the identities of other sites. However, experimental work with NP (Gong et al., 2013) and other proteins (Serrano et al., 1993; Bloom et al., 2005; Bershtein et al., 2006; Bloom et al., 2006) suggests that at an evolutionary level, sites interact mostly through generic effects on stability and folding. Furthermore, the effects of mutations on stability and folding tend to be conserved during evolution (Ashenberg et al., 2013; Serrano et al., 1993). So one justification for assuming site-specific but site-independent preferences is that selection on a mutation is mostly determined by whether the protein can tolerate its effect on stability or folding, so stabilizing amino acids will be tolerated in most genetic backgrounds while destabilizing amino acids will only be tolerated in some backgrounds, as has been described experimentally (Gong et al., 2013) and theoretically (Bloom et al., 2007). A more pragmatic justification is that the work here builds off this assumption to create evolutionary models that are much better than existing alternatives.

Assume that the preferences are entirely at the amino-acid level and are indifferent to the specific codon (the study of preferences for synonymous codons is an interesting area for future work). Denote the preference of site *r* for amino-acid *a* as *π_r,a_*, where

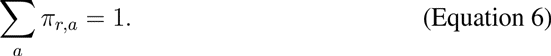

Define *π_*r,a*_/π_*r,a′*_* as the expected ratio of amino-acid *a* to *a′* after viral growth if both are initially introduced into the mutant library at equal frequency. Mutations that enhance viral growth will have larger values of *π_r,a_*, while mutations that hamper growth will have lower values of *π_r,a_*. However, *π_r,a_/π_r,a′_* cannot be simply interpreted as the fitness effect of mutating site *r* from *a* to *a′*: because most clones have multiple mutations, this ratio summarizes the effect of a mutation in a variety of related genetic backgrounds. A mutation can therefore have a ratio greater than one due to its inherent effect on viral growth or its effect on the tolerance for other mutations (or both). This analysis does not separate these factors, but experimental work (Gong et al., 2013) has shown that it is fairly common for one mutation to NP to alter the tolerance to a subsequent one.

The most naive approach is to set *π_r,a_* proportional to the frequency of amino-acid *a* in **mutvirus-p1** divided by its frequency in **mutDNA**, and then apply the normalization in Equation 6. However, such an approach is problematic for several reasons. First, it fails to account for errors (PCR, reverse-transcription) that inflate the observed frequencies of some mutations. Second, estimating ratios by dividing finite counts is notoriously statistically biased (Pearson, 1910; Ogliore et al., 2011). For example, in the limiting case where a mutation is counted once in **mutvirus-p1** and not at all in **mutDNA**, the ratio is infinity – yet in practice such low counts give little confidence that enough variants have been assayed to estimate the true effect of the mutation.

To circumvent these problems, I used an approach that explicitly accounts for the sampling statistics. The approach begins with prior estimates that the *π_r,a_* values are all equal, and that the error and mutation rates for each site are equal to the library averages. Multinomial likelihood functions give the probability of observing a set of counts given the *π_r,a_* values and the various error and mutation rates. The posterior mean of the *π_r,a_* values is estimated by MCMC.

Use the counts in **DNA** to quantify errors due to PCR and sequencing. Use the counts in **RNA** to quantify errors due to reverse transcription. Assume that transcription of the viral genes from the reverse-genetics plasmids and subsequent replication of these genes by the influenza polymerase introduces a negligible number of new mutations. The second of these assumptions is supported by the fact that the mutation frequency in **virus-p1** is close to that in **RNA** (Figure 3). The first of these assumptions is supported by the fact that stop codons are no more frequent in **RNA** than in **virus-p1** (Figure 3) – deleterious stop codons arising during transcription will be purged during viral growth, while those arising from reverse-transcription and sequencing errors will not.

At each site *r*, there are *n*_codon_ codons, indexed by *i* = 1, 2, … *n*_codon_. Let wt(*r*) denote the wildtype codon at site *r*. Let 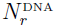 be the total number of sequencing reads at site *r* in **DNA**, and let 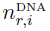 be the number of these reads that report codon *i* at site *r*, so that 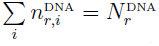. Similarly, let 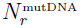, 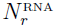, and 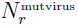 be the total number of reads at site *r* and let 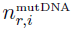, 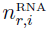, and 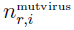 be the total number of these reads that report codon *i* at site *r* in **mutDNA**, **RNA**, and **mutvirus-p1**, respectively.

First consider the rate at which site *r* is erroneously read as some incorrect identity due to PCR or sequencing errors. Such errors are the only source of non-wildtype reads in the sequencing of **DNA**. For all *i* ≠ wt(*r*), define *ϵ_r,i_* as the rate at which site *r* is erroneously read as codon *i* in **DNA**. Define 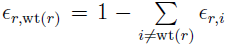 to be the rate at which site *r* is correctly read as its wildtype identity of wt (*r*) in **DNA**. Then 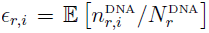 where 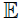 denotes the expectation value. Define 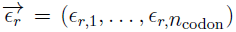 and 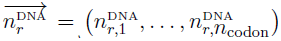 as vectors of the *ϵ_r,i_* and 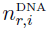 values, so the likelihood of observing 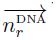 given 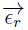 and 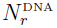 is

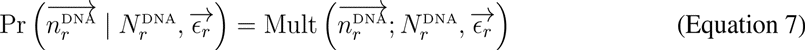

where Mult denotes the multinomial distribution.

Next consider the rate at which site *r* is erroneously copied during reverse transcription. These reverse-transcription errors combine with the PCR / sequencing errors defined by 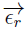 to create non-wildtype reads in **RNA**. For all *i ≠* wt (*r*), define *ρ_r,i_* as the rate at which site *r* is miscopied to *i* during reverse transcription. Define 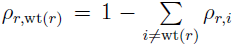 as the rate at which site *r* is correctly reverse transcribed. Ignore as negligibly rare the possibility that a site is subject to both a reverse-transcription and sequencing / PCR error within the same clone (a reasonable assumption as both *ϵ_r,i_* and *ρ*_*r*;i_ are very small for *i* ≠ wt (*r*)). Then 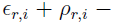 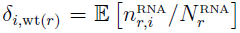 where *δ*_*i*,wt(*r*)_ is the Kronecker delta (equal to one if *i* = wt(*r*) and zero otherwise). The likelihood of observing 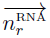 given 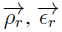, and *N*^r^_RNA_ is

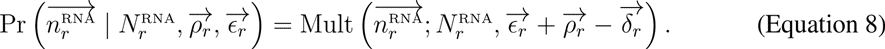

where 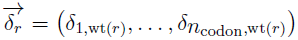 is a vector that is all zeros except for the element wt (*r*).

Next consider the rate at which site *r* is mutated to some other codon in the plasmid mutant library. These mutations combine with the PCR/ sequencing errors defined by 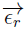 to create non-wildtype reads in **mutDNA**. For all *i* ≠ wt (*r*), define *µ_r;i_* as the rate at which site *r* is mutated to codon *i* in the mutant library. Define 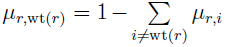 as the rate at which site *r* is not mutated. Ignore as negligibly rare the possibility that a site is subject to both a mutation and a sequencing / PCR error within the same clone. Then 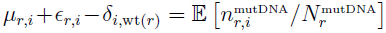. The likelihood of observing 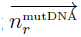 given 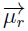, 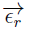, and 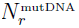 is

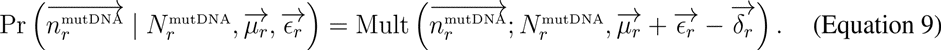

Finally, consider the effect of the preferences of each site *r* for different amino acids, as denoted by the *π_r,a_* values. Selection due to these preferences is manifested in **mutvirus**. This selection acts on the mutations in the mutant library (**µ*_r,i_*), although the actual counts in **mutvirus** are also affected by the sequencing / PCR errors (*ϵ_r,i_*) and reverse-transcription errors (*ρ_r,i_*). Again ignore as negligibly rare the possibility that a site is subject to more than one of these sources of mutation and error within a single clone. Let 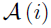 denote the amino acid encoded by codon *i*. Let 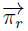 be the vector of *π_r,a_* values. Define the vector-valued function 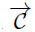 as

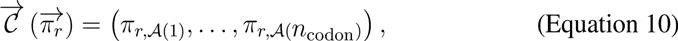

so that this function returns a *n*_codon_-element vector constructed from 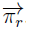. Because the selection in **mutvirus** due to the preferences 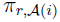 occurs after the mutagenesis *µ_r,i_* but before the reverse-transcription errors *ρ_r,i_* and the sequencing / PCR errors *ϵ_r,i_*, then 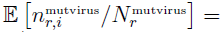 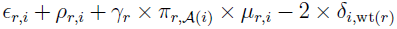 where 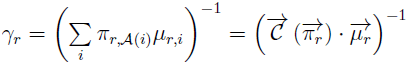 (where *·* denotes the dot product) is a normalization factor that accounts for the fact that changes in the frequency of one variant due to selection will influence the observed frequency of other variants. The likelihood of observing 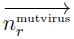 given 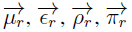, and 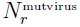 is therefore

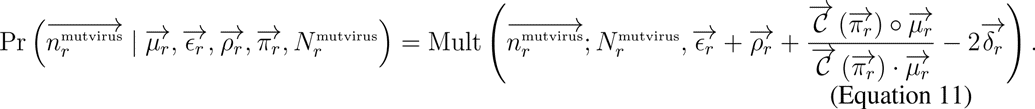

where 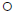 is the Hadamard (entry-wise) product.

Specify priors over 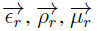, and 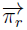 in the form of Dirichlet distributions (denoted here by Dir). For the priors over the mutation rates 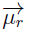, I choose Dirichlet-distribution parameters such that the mean of the prior expectation for the mutation rate at each site *r* and codon *i* is equal to the average value for all sites, estimated as the frequency in **mutDNA** minus the frequency in **DNA** (Figure 3), denoted by 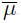. So the prior is

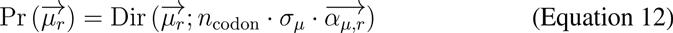

where 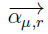 is the *n*_codon_-element vector with elements 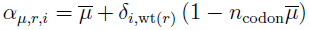 and *σ_µ_* is the scalar concentration parameter.

For the priors over *ϵ_r,i_* and *ρ_r,i_*, the Dirichlet-distribution parameters again represent the average value for all sites, but now also depend on the number of nucleotide changes in the codon mutation since sequencing / PCR and reverse-transcription errors are far more likely to lead to single-nucleotide codon changes than multiple-nucleotide codon changes (Figure 3). Let *M* (wt (*r*) *, i*) be the number of nucleotide changes in the mutation from codon wt (*r*) to codon *i*. For example, *M*(`GCA, ACA) = 1` and *M*(`GCA, ATA) = 2`. Let 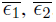, and 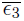 be the average error rates for one-, two-, and three-nucleotide codon mutations, respectively – these are estimated as the frequencies in **DNA**. So the prior is

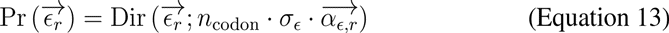

where 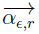 is the *n*_codon_-element vector with elements 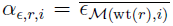 where 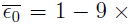 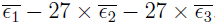, and where *σ_ϵ_* is the scalar concentration parameter.

Similarly, let 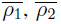, and 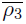 be the average reverse-transcription error rates for one-, two-, and three-nucleotide codon mutations, respectively – these are estimated as the frequencies in **RNA** minus those in **DNA**. So the prior is

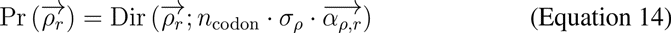

where 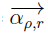 is the *n*_codon_-element vector with elements 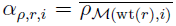 where 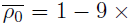 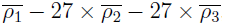, and where *σ_ρ_* is the scalar concentration parameter.

Specify a symmetric Dirichlet-distribution prior over 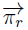 (note that any other prior, such as one that favored wildtype, would implicitly favor certain identities based empirically on the wildtype sequence, and so would not be in the spirit of the parameter-free derivation of the *π_r,a_* values employed here). Specifically, use a prior of

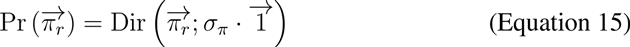

where 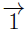 is the *n*_aa_-element vector that is all ones, and *σ_π_* is the scalar concentration parameter.

It is now possible to writ expressions for the likelihoods and posterior probabilities. Let 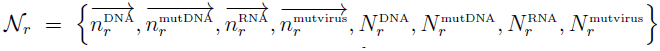 denote the full set of counts for site *r*. The likelihood of 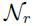 given values for the preferences and mutation/error rates is

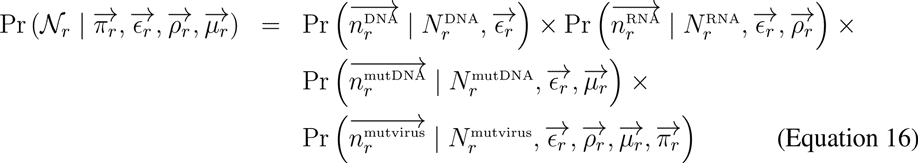

where the likelihoods that compose Equation 16 are defined by Equations Equation 7, Equation 8, Equation 9, and Equation 11. The posterior probability of a specific value for the preferences and mutation / error rates is

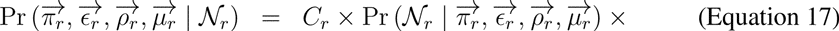

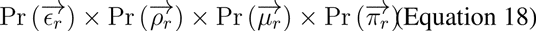

where *C_r_* is a normalization constant that does not need to be explicitly calculated in the MCMC approach used here. The posterior over the preferences 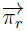 can be calculated by integrating over Equation 17 to give

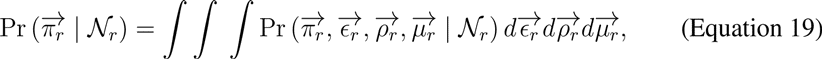

where the integration is performed by MCMC. The posterior is summarized by its mean,

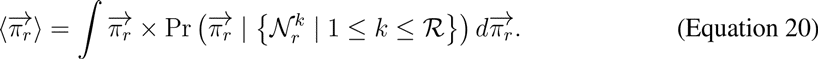

In practice, each replicate consists of four libraries (WT-1, WT-2, N334H-1, and N334H-2) – the posterior mean preferences inferred for each library within a replicate are averaged to give the estimated preferences for that replicate. The preferences within each replicate are highly correlated regardless of whether **mutvirus-p1** or **mutvirus-p2** is used as the **mutvirus** data set (Figure 5A,B). This correlation between passages is consistent with the interpretation of the preferences as the fraction of genetic backgrounds that tolerate a mutation (if it was a selection coefficient, there should be further enrichment upon further passage). The preferences averaged over both replicates serve as the “best” estimate, and are displayed in Figure 5D. This figure was created using the WebLogo 3 program (Schneider and Stephens, 1990; Crooks et al., 2004).

Figure 5D also shows relative solvent accessibility (RSA) and secondary structure for residues present in chain C of NP crystal structure PDB 2IQH (Ye et al., 2006). The total accessible surface area (ASA) and the secondary structure for each residue in this monomer alone was calculated using DSSP (Kabsch and Sander, 1983; Joosten et al., 2011). The RSAs are the total ASA divided by the maximum ASA defined in (Tien et al., 2013). The secondary structure codes returned by DSSP were grouped into three classes: helix (DSSP codes G, H, or I), strand (DSSP codes B or E), and loop (any other DSSP code).

### Phylogenetic analyses

A set of NP coding sequences was assembled for human influenza lineages descended from a close relative the 1918 virus (H1N1 from 1918 to 1957, H2N2 from 1957 to 1968, H3N2 from 1968 to 2013, and seasonal H1N1 from 1977 to 2008). All full-length NP sequences from the Influenza Virus Resource (Bao et al., 2008) were downloaded, and up to three unique sequences per year from each of the four lineages described above were retained. These sequences were aligned using EMBOSS needle (Rice et al., 2000). Outlier sequences that correspond to heavily lab-adapted strains, lab recombinants, mis-annotated sequences, or zoonotic transfers (for example, a small number of human H3N2 strains are from zoonotic swine variant H3N2 rather than the main human H3N2 lineage) were removed. This was done by first removing known outliers in the influenza databases (Krasnitz et al., 2008), and then using an analysis with RAxML (Stamatakis, 2006) and Path-O-Gen (http://tree.bio.ed.ac.uk/software/pathogen/) to remove remaining sequences that were extreme outliers from the molecular clock. The final alignment after removing outliers consisted of 274 unique NP sequences.

Maximum-likelihood phylogenetic trees were constructed using *codonPhyML* (Gil et al., 2013). Two substitution models were used. The first was *GY94* (Goldman and Yang, 1994) using *CF3×4* equilibrium frequencies (Pond et al., 2010), a single transition-transversion ratio optimized by maximum likelihood, and a synonymous-nonsynonymous ratio drawn from four discrete gamma-distributed categories with mean and shape parameter optimized by maximum likelihood (Yang et al., 2000). The second was *KOSI07*+*F* (Kosiol et al., 2007), optimizing the relative transversion-transition ratio by maximum likelihood, and letting the relative synonymous-nonsynonymous ratio again be drawn from four gamma-distributed categories with mean and shape parameter optimized by maximum likelihood. The trees produced by *codonPhyML* are unrooted. These trees were rooted using Path-O-Gen (http://tree.bio.ed.ac.uk/software/pathogen/), and visualized with FigTree (http://tree.bio.ed.ac.uk/software/figtree/) to create the images in Figure 7. The tree topologies are extremely similar for both models.

The evolutionary models were compared by using them to optimize the branch lengths of the fixed tree topologies in Figure 7 so as to maximize the likelihood using *HYPHY* (Pond et al., 2005) for sites 2 to 498 (site 1 was not included, since the N-terminal methionine is conserved and was not mutated in the plasmid mutant libraries). *HYPHY* was used to calculate all likelihoods (even for models that could be handled by *codonPhyML*) for consistency in case these programs differ slightly in numerical accuracy. The results are shown in Table 7. Regardless of which tree topology was used, the experimentally determined evolutionary models outperformed all variants of *GY94* and *KOSI07*+*F*. The experimentally determined evolutionary models performed best when using the preferences determined from the combined data from both replicates and using Equation 3 to compute the fixation probabilities. Using the data from just one replicate also outperforms *GY94* and *KOSI07*+*F*, although the likelihoods are slightly worse. In terms of the completeness with which mutations are sampled in the mutant viruses, *replicate A* is superior to *replicate B* as discussed above – and the former replicate gives higher likelihoods. If the fixation probabilities are instead determined using the method of Halpern and Bruno (Halpern and Bruno, 1998) as in Equation 4, the experimentally determined models still outperform *GY94* and *KOSI07*+*F* – but the likelihoods are substantially worse. To check that the experimentally determined models really do utilize the site-specific preferences information, the preferences were randomized among sites and likelihoods were computed. These randomized models perform vastly worse than any of the alternatives.

The variants of *GY94* and *KOSI07*+*F* tested are listed in Table 6. Various methods were used to estimate the nonsynonymous-synonymous ratio (*ω*): a single *ω* optimized by maximum likelihood; three discrete categories of *ω* < 1, *ω* = 1, and *ω* > 1 with the proportions and the *ω* values ≠ 1 estimated by maximum likelihood; *ω* drawn from four gamma-distributed categories with mean and shape estimated by maximum likelihood; and a beta distribution (10 categories) plus an additional category of *ω* > 1 with the shape parameters, *ω* > 1 value, and proportion in the final category estimated by maximum likelihood. These models are referred to M0, M2a, M5, and M7 in the literature (Yang et al., 2005). Another model optimized a different *ω* for each branch. Another model optimized a single *ω* but allowed the rates to be drawn from four gamma-distributed categories. Parameters were counted as follows: all contained equilibrium frequency parameters that were empirically estimated from the sequences under analysis: there are 9 such parameters for *GY94* using *CF3x4* (Goldman and Yang, 1994; Pond et al., 2010) and 60 such parameters for *KOSI07*+*F* (Kosiol et al., 2007). In addition, all variants contain a transition-transversion ratio optimized by likelihood. Finally, all variants contain one or more *ω* parameters as described above.

## Acknowledgments

Thanks to D. Fowler, J. Kitzman, A. Adey, O. Ashenberg, and T. Bedford for helpful discussions. This work was supported by the National Institute of General Medical Sciences of the National Institutes of Health (grant number R01 GM102198).

---

**Supplementary figure 1:**
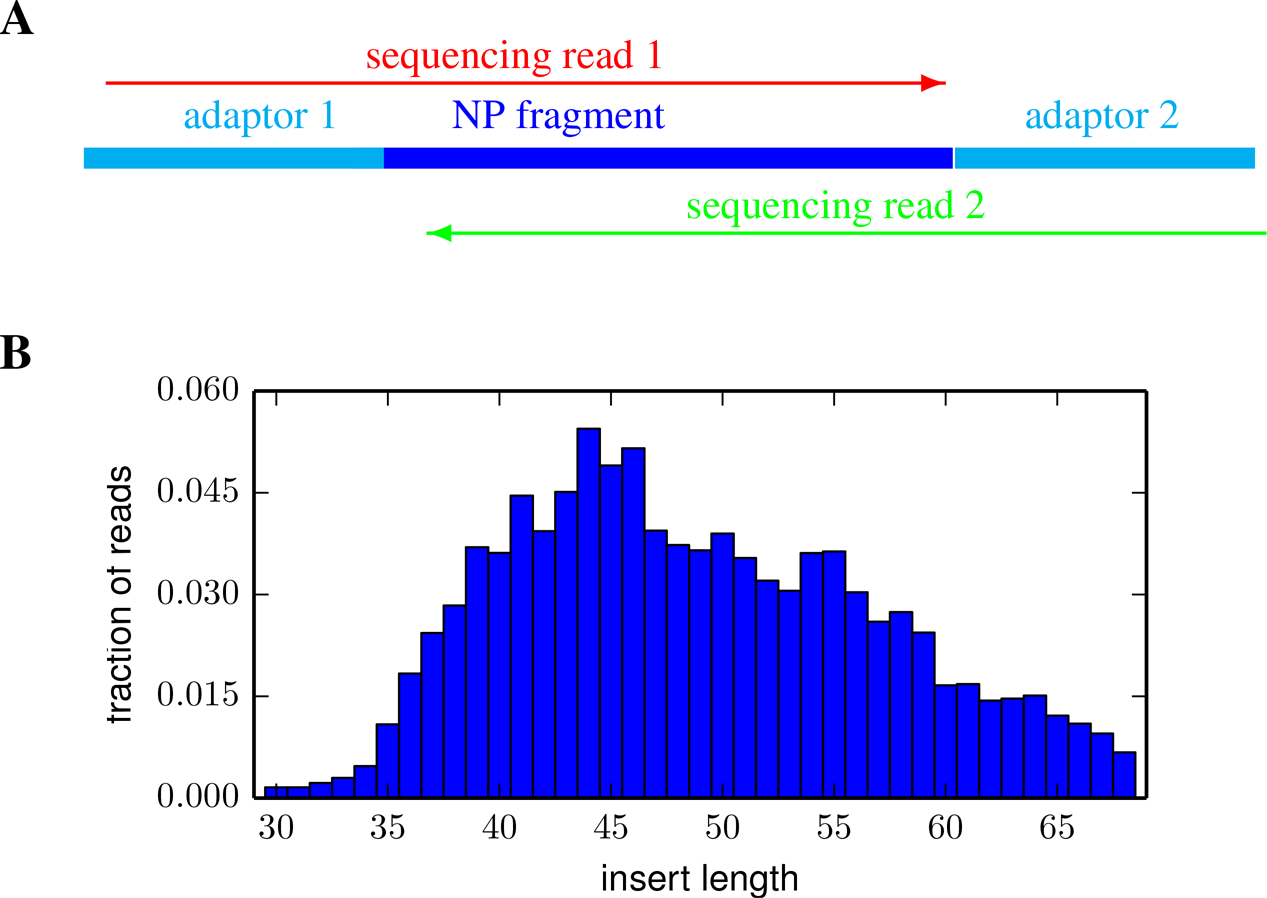
Illumina sequencing accuracy was increased by using overlapping paired-end reads. **(A)** The strategy is to shear the NP fragments to about 50 nucleotides in length, and then sequence with overlapping paired-end reads. This provides double coverage, and only codon identities for which both reads agree are called. **(B)** The actual length distribution of alignable paired reads that had at least 30 nucleotides of overlap (lengths between 30 and 70 nucleotides) for a typical sample.

**Supplementary figure 2:**
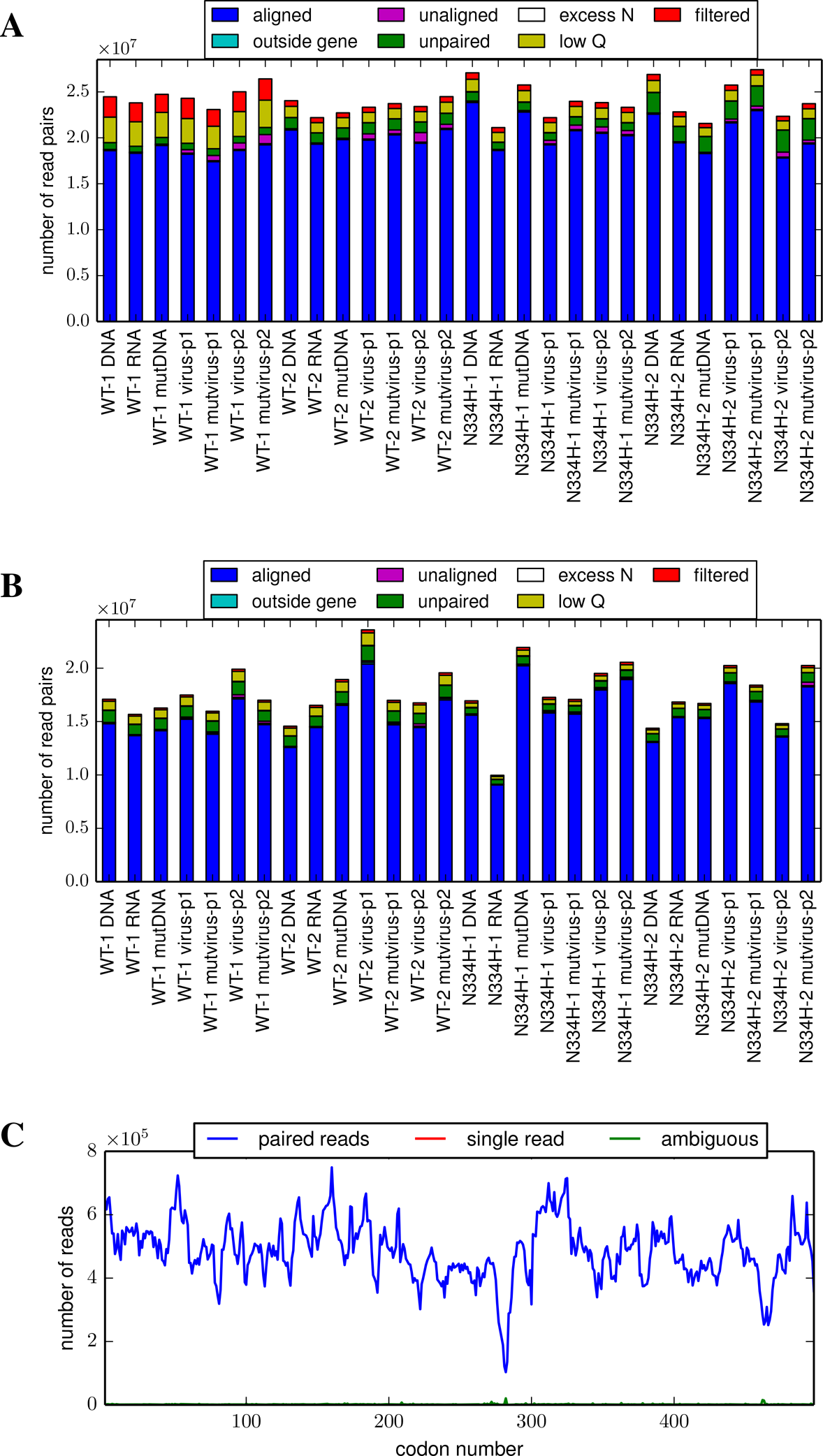
The number of alignable paired reads for each sample from **(A)** *replicate A* and **(B)** *replicate B*. Most read pairs passed the various filters, could be overlapped, and could have their overlap aligned to NP. **(C)** The read depth across the primary sequence for a typical sample. The read depth was not entirely uniform along the primary sequence, probably due to biases in fragmentation locations. The computer code used to generate these figures is available at http://jbloom.github.io/mapmuts/example_2013Analysis_Influenza_NP_Aichi68.html.

**Supplementary file 1:** The file *Preferences.xlsx* is an Excel table of the inferred amino-acid preferences at each site from the combined passage 1 data for the two replicates.

**Supplementary file 2:** The file *EvolutionaryEquilibriumFreqs.xlsx* is an Excel table of the expected equilibrium frequencies of the amino acids during evolution.

